# Eight principal chromatin states functionally segregate the fly genome into developmental and housekeeping roles

**DOI:** 10.1101/2022.10.30.514435

**Authors:** Caroline Delandre, John P. D. McMullen, Jonas Paulsen, Philippe Collas, Owen J. Marshall

## Abstract

Different chromatin forms, or states, represent a fundamental means of controlling gene regulation. Chromatin states have been studied through either the distribution of histone modifications (e.g. ^1–5^) or more rarely via the occupancy of chromatin proteins ^6–8^. However, these two approaches disagree on the nature and composition of active chromatin states ^2,9^ and modelling chromatin via both histone marks and chromatin proteins has been lacking. Here, combining protein and histone mark profiles, we show that chromatin in *Drosophila melanogaster* is organised into eight principle chromatin states that have consistent forms and constituents across cell types. These states form through the association of the Swi/Snf chromatin remodelling complex, Polycomb Group (PcG)/H3K27me3, HP1a/H3K9me3 or H3K36me3 complexes with either active complexes (RNA Pol/COMPASS/H3K4me3/NuRF) or repressive marks (histone H1 and nuclear lamin occupancy). Enhancers, core promoters, transcription factor motifs, and gene bodies show distinct chromatin state preferences that separate by developmental and housekeeping/metabolic gene ontology. Within the 3D genome, chromatin states add an additional level of compartmentalisation through self-association of topologically associated domains (TADs) of the same state. Our results suggest that the epigenetic landscape is organised by the binding of chromatin remodellers and repressive complexes, and that through chromatin states the genome is fundamentally segregated into developmental and housekeeping/metabolic roles.

## Eight principal chromatin states organise the fly genome

To understand the combined organisation of chromatin proteins and histone marks, we developed a new software package for modelling chromatin states. Our system, which we term ChroMATIC (<u>Chro</u>matin Hidden <u>M</u>arkov Models, <u>A</u>IC-weigh<u>t</u>ed, <u>i</u>nferred and <u>c</u>lustered), allows for the detection of variable levels of protein binding and histone mark occupancy via multivariate Gaussian Hidden Markov Models (HMMs), and applies the power of multi-model inference to chromatin state modelling, illustrating models as both state emission heatmaps and state transition network graphs (Fig. 1a; see Methods for details).

**Figure 1:**
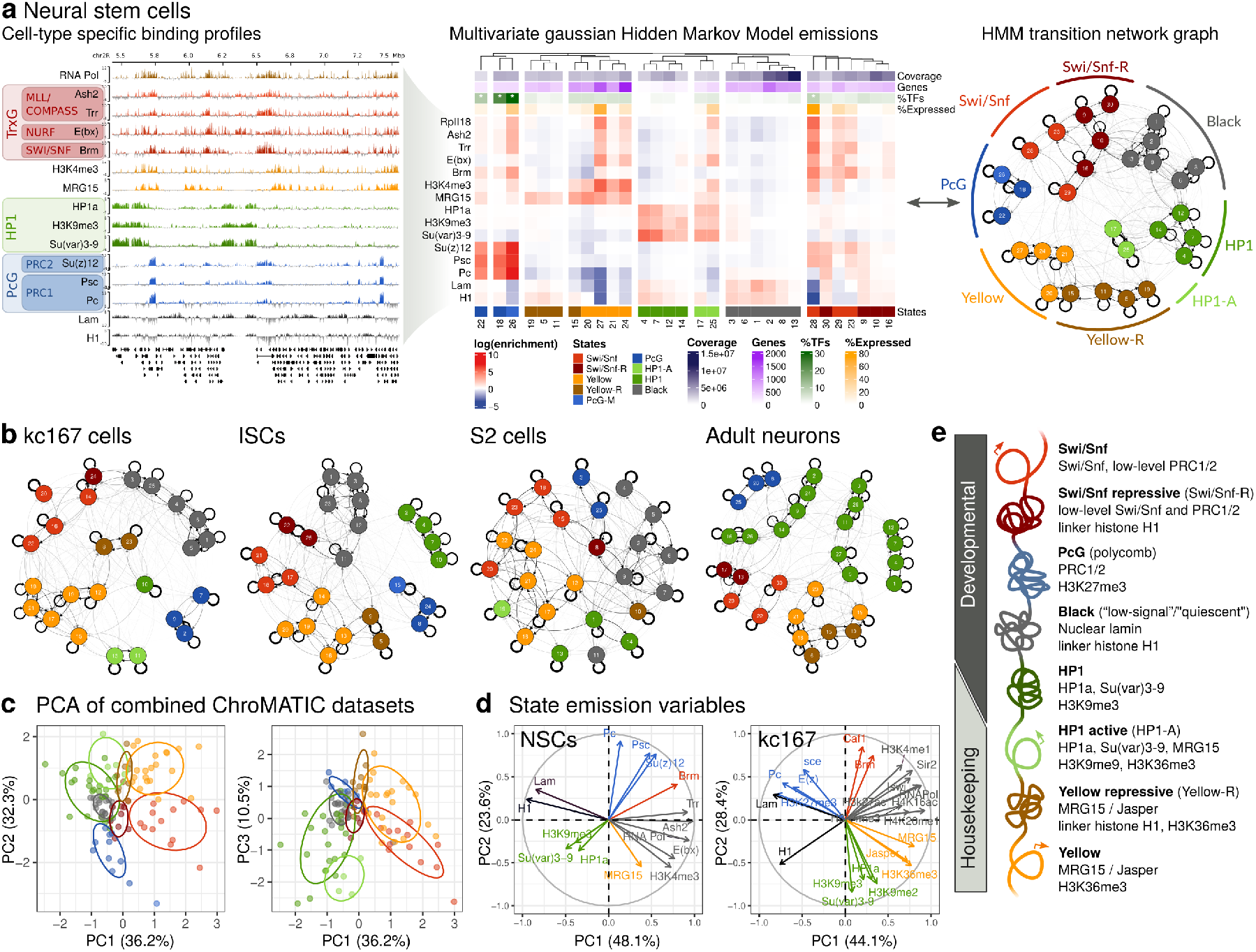
Chromatin states cluster into eight principle types in *Drosophila*. (a) Targeted DamID profiles for 13 proteins and 2 histone modification marks were obtained from neural stem cells (NSCs); larger chromatin complex groupings are illustrated. These profiles were used to generate multivariate gaussian HMMs via ChroMATIC; the best-fitting model is illustrated as both a state emissions heatmap and state transitions network graph. In the heatmaps, genomic coverage of each state, the number of genes covered, the number expressed (determined via RNA polymerase occupancy) and the percentage of genes within a state that are transcription factors (TFs; * = *P*<0.01, Benjamini-Hochberg adjusted, Fisher’s exact test) are also illustrated. (b) State transitions as determined through ChroMATIC modelling for three additional dividing cell types (kc167 cells, intestinal stem cells (ISCs) and S2 cells) and a terminally-differentiated cell type (adult mature neurons). (c) PCA of shared proteins emission means and relative transition network graph distances, formed from all datasets except ISCs; points and confidence ellipses are coloured by chromatin state. (d) Variable correlation plots from PCA of ChroMATIC model emissions; distinguishing marks and proteins have been coloured by groupings. (e) A schematic of the eight chromatin states identified in this study with characteristic chromatin proteins and histone marks listed for each.

We applied ChroMATIC to chromatin datasets from five separate *Drosophila* cell types (Table S1), covering major chromatin remodellers, activating and repressive protein complexes, commonly-profiled histone modifications and RNA polymerase (Table S2). Via Targeted DamID (TaDa) and a new technique (“ChromaTaDa”; Fig. S1) we profiled 13 chromatin proteins and 2 histone marks in larval neural stem cells (NSCs), and 9 chromatin proteins in mature adult neurons (Fig. S3). We also reprocessed existing DamID and ChIP-seq datasets from the early embryonic cell line kc167 (14 proteins + 9 histone marks; Fig. S4), S2 cells (9 proteins + 7 histone marks; Fig. S5) and Intestinal Stem Cells (ISCs) (5 proteins; Fig. S6) (Fig. 1b).

Our chromatin models revealed a separation of the genome into eight broad chromatin states (Fig. 1a,b) which we could show to be equivalent across cell-types via principal component analysis (PCA) of the combined HMM state emissions (Figs 1c). Clustering via state transitions (Figs. 1a,S4,S3,S5,S6) and PCA of state emissions by cell type (Figs. 1d, S2) indicated that these states formed through the association of the Swi/Snf, HP1a/H3K9me3, PcG/H3K27me3 or H3K36me3 complexes with either active complexes (RNA Pol/COMPASS/H3K4me3/NuRF) or repressive marks (histone H1 and nuclear lamin occupancy) (Fig. 1e). These constituents have established interactions that suggest a physical basis for chromatin states ^10–17^, most notably the refractory relationships between PcG/H3K27me3 and H3K36me3^15^, and between Swi/Snf and H3K9me3/HP1 heterochromatin ^12^.

These eight chromatin states shared three repressive states in common with earlier protein ^6^ and histone mark studies ^2,4^: Polycomb Group (PcG) facultative heterochromatin, HP1 constitutive heterochromatin, and a silent “Black” (or “low-signal”) chromatin state. This latter state was enriched for the linker histone H1 and nuclear lamin occupancy, devoid of other profiled proteins or histone marks, and shared similar genomic coverage with earlier classifications of both Black chromatin in kc167 cells ^6^ (Fig. S7a) and a low-signal state in S2 cells ^2^ (Fig. S7b).

However, in contrast to previous chromatin state modelling, regions of the genome with Swi/Snf, MRG15/H3K36me3 or HP1a/H3K9me3 occupancy unexpectedly separated into both active (Swi/Snf, “Yellow” ^6^ and HP1-Active (“HP1-A”)) and silent (Swi/Snf-Repressive (“Swi/Snf-R”), Yellow-Repressive (“Yellow-R”) and HP1) forms. Assigning each gene a chromatin state by modal coverage, we found this separation into active and repressive states to match existing gene expression data (Fig. 2a). We have previously reported the presence of a silent Swi/Snf-enriched chromatin state in neural development ^8^ – an observation that fits with a body of work noting a repressive function for Brm and the Swi/Snf complex ^11,18–20^. Transcriptionally-active HP1a/H3K9me3 chromatin has been previously described at fly heterochromatic regions ^14,21^ and in the euchromatic regions of differentiated human fibroblasts ^22^; here, we also observe this signature within the euchromatin of stem cells, implying that this means of gene activation is common across cell types and not restricted to pericentric heterochromatin. Yellow-R, a transcriptionally-silent form of MRG15/H3K36me3 chromatin, has not been previously described to our knowledge.

**Figure 2:**
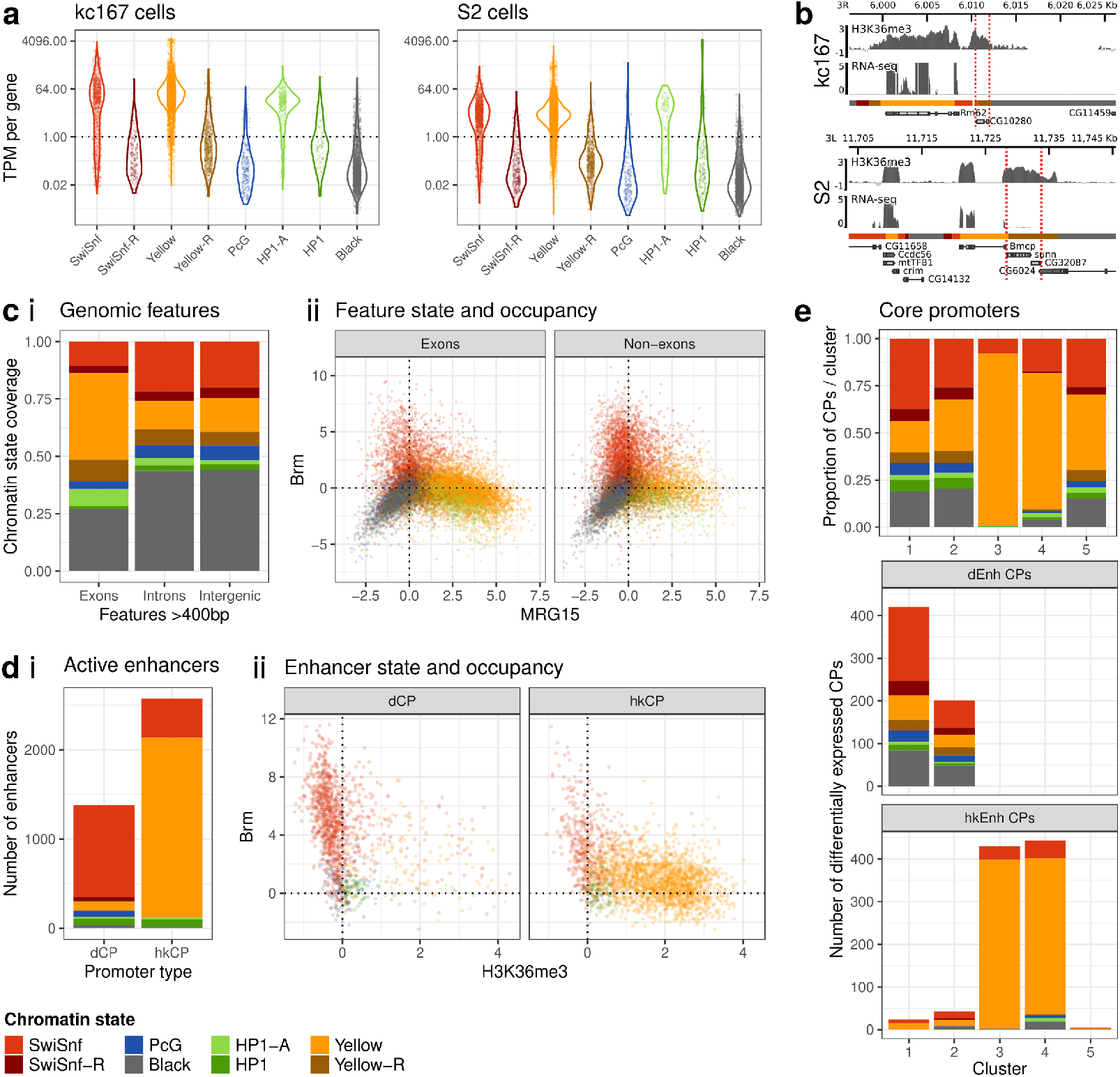
Chromatin states divide genomic features into developmental and housekeeping roles. (a) RNA-seq expression levels of genes by chromatin state for kc167 and S2 cell lines illustrates the active and repressive nature of the eight chromatin states. (b) Yellow-R regions with high H3K36me3 are transcriptionally silent. Example genome plots showing H3K36me3 occupancy, RNA-seq pseudoalignments, and chromatin states within the genome for kc167 and S2 cells; Yellow-R regions covering genes are highlighted with dashed lines. (c) Chromatin states in kc167 cells are distributed across exons, introns and intergenic regions (i) and separated by state on key chromatin protein occupancy (ii). (d) Active enhancer regions identified by STARR-seq in S2 cells, coloured by chromatin state, show clear separation of chromatin state occupancy by core promoter type in S2 cells (i), with Yellow chromatin enriched over active housekeeping enhancers, and Swi/Snf chromatin enriched over active developmental proteins; and (ii) differing levels of Brm and H3K36me3 occupancy by enhancer class. (e) Promoter (CP) clusters identified by STAP-seq show biases in chromatin state genomic occupancy. For core promoters that were differentially activated by either a developmental enhancer (dEnh) or a housekeeping enhancer (hkEnh) in S2 cells, the chromatin state of the four active core promoter clusters strongly predicted enhancer-core-promoter ontology.

The presence of silent Yellow-R chromatin was surprising given earlier studies linking H3K36me3 with actively transcribed exons ^23^. However, clear instances of H3K36me3-enriched but transcriptionally-silent loci were present throughout the genome (Fig. 2b), chromatin states other than Yellow/Yellow-R also covered exons (Fig. 2c (i)), and there was substantial coverage of Yellow and Yellow-R chromatin over introns and intergenic regions (Fig. 2c (i)) linked with MRG15/H3K36me3 occupancy (Figs 2c(ii),S8). H3K36me3 chromatin is thus not representative of exon features or transcription *per se*.

In neural stem cells and intestinal stem cells – both of which can give rise to multiple downstream cell types and sub-lineages – we also observed a transcriptionally-active signature within some of the PcG states. We have previously shown that this “PcG-mixed” signature in NSCs covers TFs that specify neuronal subtypes, and derives from a heterogeneous population of cells with either active or repressive states at these loci ^8^. In concordance with this, we failed to detect an active PcG signature in either the S2 (Fig. S5) or kc167 (Fig. S4) homogeneous cultured cell lines.

Our modelling reveals a new level of complexity in chromatin state organisation, in which marks of active transcription and condensed chromatin are required to understand regulatory context. We note that while histone marks and chromatin-protein complexes overlap for many states, none of the commonly-profiled histone marks in this study associated with the Swi/Snf complex. This may in part explain discrepancies between previous protein-based and histone-mark-based modelling studies, and underlines the importance of profiling chromatin proteins when modelling chromatin states.

### Chromatin states separate enhancers and core promoters into developmental or house-keeping roles

*Drosophila* enhancers have been shown to separate into developmental and housekeeping roles ^24^, and we wondered whether these two categories of enhancers associated with different chromatin states. Assigning chromatin states to enhancers that activated developmental or housekeeping core promoters (CPs) in S2 cells ^24^, we observed that developmental enhancers were significantly depleted for Yellow chromatin and H3K36me3, and enriched for Swi/Snf chromatin and Brm occupancy, with the situation reversed for housekeeping enhancers (Fig. 2d; Fig. S9). Profiling the chromatin states of these same enhancers in kc167 cells and NSCs, we observed a similar result, albeit with some S2 cell developmental enhancers covered in repressive chromatin marks in NSCs (Fig. S10).

Given the link between enhancers and core promoters, we next looked at the five distinct CP classes identified in S2 cells via transcription cofactor binding ^25^ (Fig. S11). Assigning chromatin states to CPs, we observed a significant depletion of Yellow chromatin in two CP types, and a significant enrichment over a further two types (Fig. 2e; Fig. S12). This divergence became even more pronounced on CPs that were differentially activated by a developmental or house-keeping enhancer (Fig. 2e). Combined, these data suggest that chromatin states fundamentally separate enhancer and core promoter regulatory elements along developmental and housekeeping lines. Importantly, our chromatin state models allow us to relate enhancers and core promoters identified from ectopic, transgenic constructs to the chromatin contexts at their native locus, demonstrating that regulatory links between enhancers and core promoters translate to differing chromatin states *in vivo*.

These data fit well with recent findings that the Swi/Snf remodelling complex is bound at and required for developmental enhancer activation in S2 cells, specifically affecting developmental gene expression when depleted ^26^. This study also looked at the effect of depleting other chromatin remodeling complexes, including NuRF and INO80, finding no ontological preference for these enhancers and – in concordance with our chromatin state modelling – observing that NuRF bound indiscriminately over both classes of promoters ^26^. However, this work failed to profile the Tip60/p400 remodelling complex that also associates with the Yellow chromatin component MRG15^27^, and is tempting to speculate that this complex may be a specific remodeller for housekeeping regulatory elements.

### Transcription factor binding motifs separate by chromatin state

Given the clear separation of chromatin states with enhancer types, we wondered whether transcription factors (TFs) would also show preferences for chromatin state. Using RCisTarget ^29^ to search for significant enrichment of TF binding motifs associated with genes in different chromatin states, we observed a clear segregation of TF motifs by chromatin state (Fig. 3a). In particular, clusters of specific TF motifs were observed for Yellow and PcG chromatin states from all cell types. Motifs for the cell-growth regulating TF Cropped (crp) ^30^, the insulator BEAF-32^31^, and the polarity and proliferation regulating TF Zif ^32^ were enriched in Yellow chromatin; whereas motifs for classic cell-fate and segmentation regulators such as Kr ^33^, ems ^34^ and bowl ^35^ were enriched in PcG chromatin (Figs. 3a, S13).

**Figure 3:**
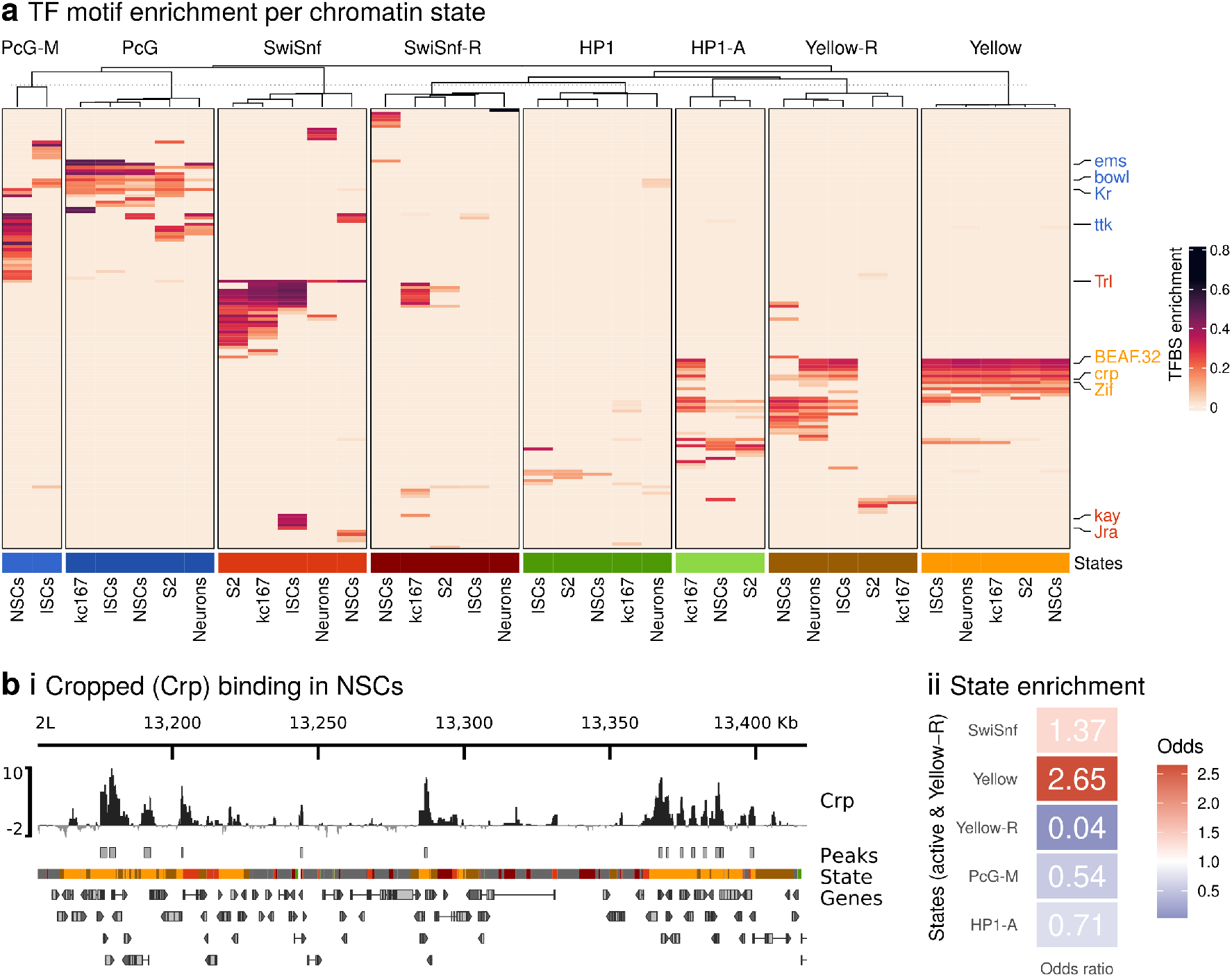
Chromatin states are differentially enriched for TF motifs across cell types. (a) For each cell type, genes were grouped by state, enriched TF motifs searched for using RcisTarget, and the proportion of genes in each state for each enriched motif recorded. High-confidence motifs were then clustered across all cell types and by chromatin state. Highlighted TF annotations are coloured by the chromatin state they are found enriched within. (b) (i) The binding of the TF Cropped (Crp) in NSCs, determined via Nanodam ^28^, is (ii) strongly and significantly enriched in Yellow, but not Yellow-R chromatin; peaks (FDR<0.01) are illustrated.

We also observed cell-type- and state-specific enrichment for some factors – such as the Jun/Fos fly orthologues Jra and Kay in active Swi/Snf chromatin in ISCs, and the neural-fate specifying TF tramtrack (ttk) ^36^ enriched in both NSC and neuronal PcG states. These data suggest that chromatin states and TF binding are intimately linked, with common conserved motifs present together with cell-type-specific factors.

To determine whether the separation of enriched TF motifs translated to state-specific TF binding *in vivo*, we profiled the binding of the Yellow-chromatin-enriched factor Cropped in NSCs (Fig. 3B(i)). Cropped has been shown to regulate cell growth and division ^30^, fitting well with a role in housekeeping gene regulation. Although the consensus motif for Cropped was enriched in both Yellow-R and Yellow chromatin in NSCs, we found that Cropped binding was significantly enriched only within active Yellow chromatin (Fig. 3B(ii)). Thus, while motif enrichment can translate to state-specific TF binding, chromatin state also plays a role in determining the final bound location.

### Yellow and Black chromatin dominate and divide the epigenetic landscape into housekeeping and developmental roles

Given the separation of transcription factor motifs, enhancers and core promoters by chromatin state, we were interested to know if genes would also separate by chromatin state and gene ontology (GO). Strikingly, clustering genes from all cell types by chromatin state revealed a developmental genome landscape dominated by active Yellow and repressive Black chromatin states (Fig. 4). In line with the association with housekeeping enhancers and core promoters described above, GO enrichment analysis revealed that Yellow chromatin associated with broadly-expressed metabolic and housekeeping genes (Fig. 4, Fig. S14). While many of these genes remained expressed in active Yellow chromatin across all studied cell types, some transitioned to active HP1-A or were repressed in Yellow-R chromatin. Conversely, genes associated with developmental processes were broadly repressed in silent Black chromatin, and transitioned to Swi/Snf and Swi/Snf-R chromatin in cell-type-specific clusters (Fig. 4, Fig. S14), matching our data for developmental enhancers. These clusters of Swi/Snf-activated genes separated by cellular type and function, with specific activation of neuroblast-associated genes in NSCs, proteolysis-associated genes in ISCs, and signalling genes in neurons (Fig. 4, Fig. S14). In contrast to dividing celltypes, terminally-differentiated neurons showed repression of many developmental gene clusters by HP1 heterochromatin.

**Figure 4:**
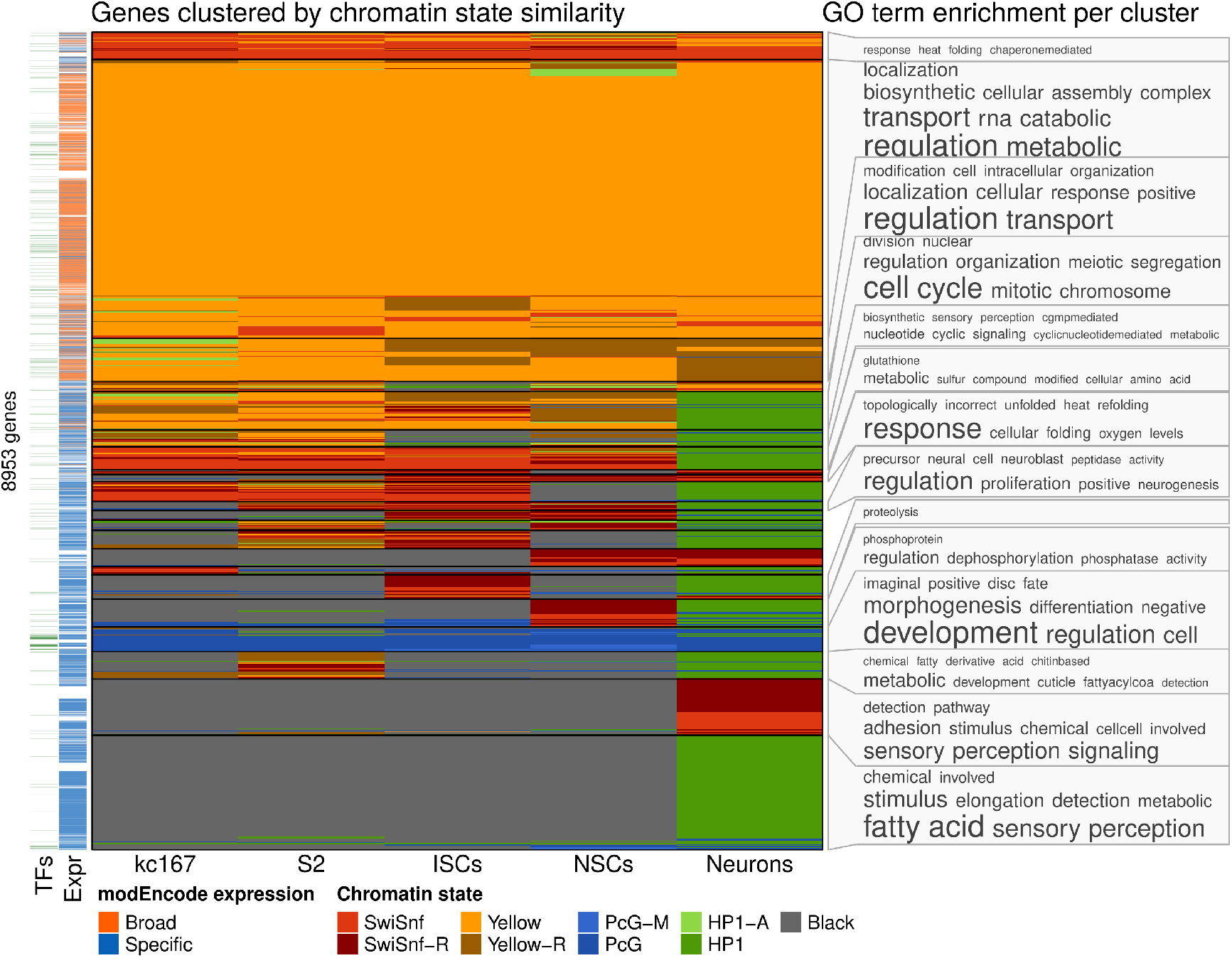
The Yellow and Black chromatin states dominate the transcriptional landscape in dividing cell types. Genes with high-confidence chromatin state calls were clustered on chromatin state. Enriched GO terms for each cluster are shown, represented as a word cloud. A classification of genes as either broadly or specifically expressed, derived from modEncode data ^4^, is shown, along with genes that encode a transcription factor ^37^.

Notably different from all other clusters, a single group of genes was present in PcG chromatin across cell types, and was highly-enriched for transcription factors with established roles in cell fate, development and differentiation (Fig. S14). In agreement with previous studies ^2,4,8,38^, we otherwise found few developmental genes regulated by PcG chromatin in any context.

### Chromatin state transitions during neurogenesis reveal unexpected roles for Yellow-R chromatin

Our new models for NSCs and neurons also allowed us to reanalyse chromatin state transitions occurring during neurogenesis. Confirming our previous findings of large-scale chromatin remodelling ^8^ (Fig. 5a), genes involved in synaptic signalling and axonogenesis transitioned from Black and Swi/Snf-R chromatin, respectively, to active Swi/Snf chromatin in neurons, while NSC identity genes transitioned from active Swi/Snf chromatin to repressive HP1 chromatin (Fig. 5b(i),c).

**Figure 5:**
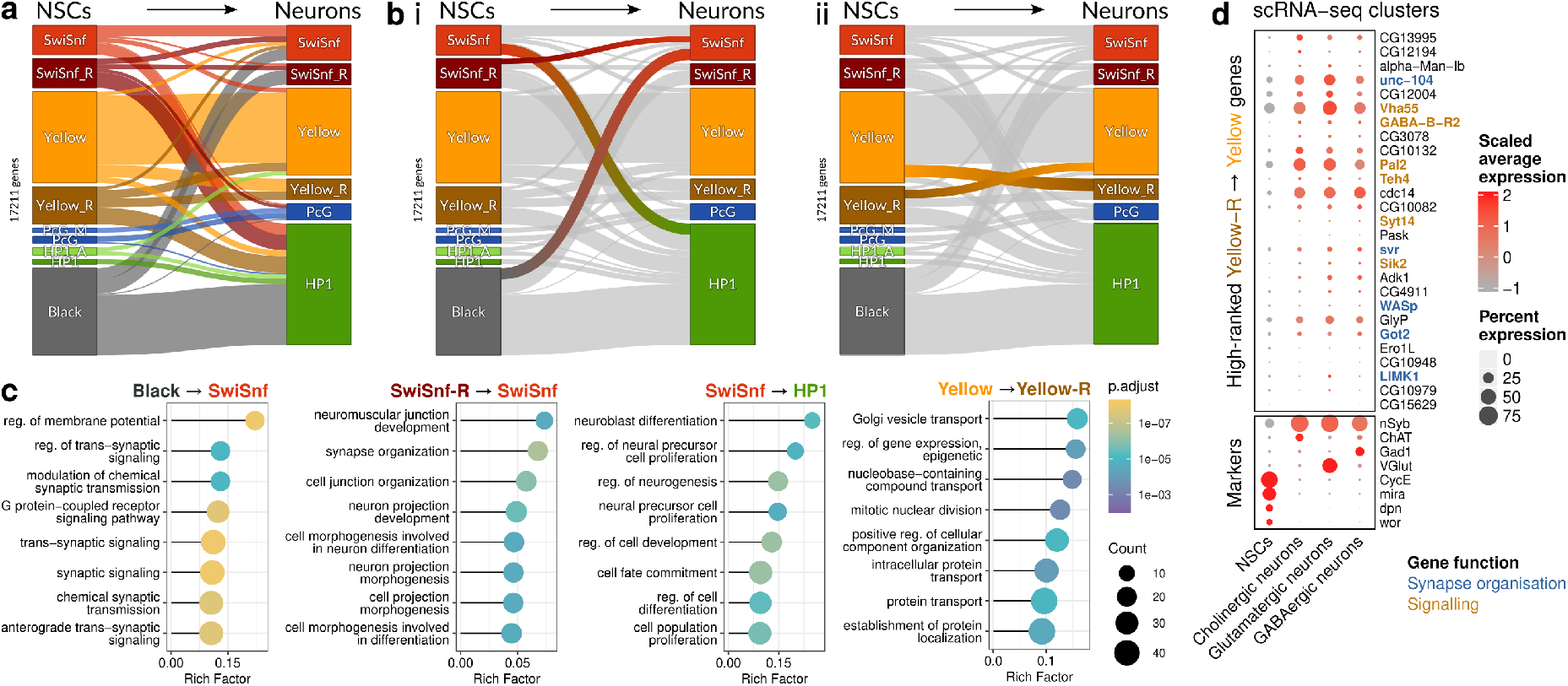
Chromatin state transitions during neural development highlights key remodelling events influencing cell fate. (a) Assigning all genes in NSCs and neurons a chromatin state based on modal state occupancy reveals extensive chromatin remodelling during differentiation, including (b) genes transitioning between active Swi/Snf and repressive states (i) and between active and repressive forms of Yellow chromatin (ii). Functional roles of different transitions are illustrated via GO term enrichments (c). Although not significantly enriched for specific ontologies, investigation of the highest-ranking genes transitioning from Yellow-R in NSCs to Yellow states in neurons revealed a number of genes involved in synaptic structure and signalling, with scRNA-seq data (reanalysed from ^39^) confirming the change in expression status between cell types (d).

However, we also observed genes that transitioned between the active Yellow and silent Yellow-R states, and *vice versa* (Fig. 5b(ii)). Genes repressed by Yellow-R in neurons fell under broad cellular transport and mitotic gene ontology categories, indicating that not all genes involved in cell division are repressed by HP1 heterochromatin upon terminal differentiation (Fig. 5c). Although not significantly enriched for ontology, genes repressed by Yellow-R chromatin in NSCs and activated by Yellow chromatin in neurons included critical neuronal genes involved in assembling and maintaining neuronal synapses (e.g. *Got2* ^40^, *unc-104* ^41^) and synaptic signalling *(e*.*g. Pal2* ^*42*^, *GABA-B-R2*^*43*^). Reanalysis of recent scRNA-seq data for the *Drosophila* larval brain ^39^ confirmed that these MRG15-enriched genes were silent in NSCs and broadly expressed across mature neuron subtypes (Fig. 5d). Combined, these data suggest that Yellow chromatin transitions are an important and previously unrecognised means of controlling gene expression during changes in cell fate.

### Chromatin states compartmentalise the 3D genome

Chromatin state domain boundaries have been linked with topologically-associated domains (TADs) ^44^, and TADs associate into A/B compartments based on transcriptional status ^45^. These compartments can be further divided into sub-compartments, partially separated by histone modifications ^46^, and we wondered whether TADs of similar chromatin state would also exhibit longrange associations within the 3D genome. Reprocessing existing HiC ^47^ and Lamin B binding data ^6^ (Fig. 6a), we adapted Chrom3D ^48,49^to model the 3D genome of kc167 cells (Fig. 6b(i)). Chrom3D uses TADs as its fundamental modelling unit, relying upon statistically-significant inter-TAD interactions and TAD-Lamina interactions to infer 3D genome structure. Assigning each TAD a chromatin state via modal chromatin state coverage thus allowed us to generate 3D genome models of chromatin states, illustrating known nuclear features such as the clustering of pericentric heterochromatin regions into a single structure termed the chromocentre (Fig. 6b(ii)). Using the significantly-enriched inter-TAD associations called via Chrom3D, we then looked at whether chromatin states associated on a TAD level within the 3D genome. Although Yellow-R TADs were not frequent enough to call enrichment for (reflecting the overall low extent of Yellow-R chromatin in kc167 cells), for all other states we observed a significantly-enriched self-association of chromatin states (Fig. 6c,d; Fig. S15). We also found weaker, but still significantly-enriched associations between TADs of related pairs of states, including associations between Swi/Snf and Swi/Snf-R, HP1 and HP1-A, and HP1-A and Yellow states (Fig. 6c,d). There has been increasing evidence of liquid-liquid phase-separation of chromatin states ^50–52^, including HP1 heterochromatin ^50^ and PcG chromatin ^52^. These data, showing long-range preferential associations between TADs of the same chromatin state, and between states with similar protein binding, provide further support to a phase separation model.

**Figure 6:**
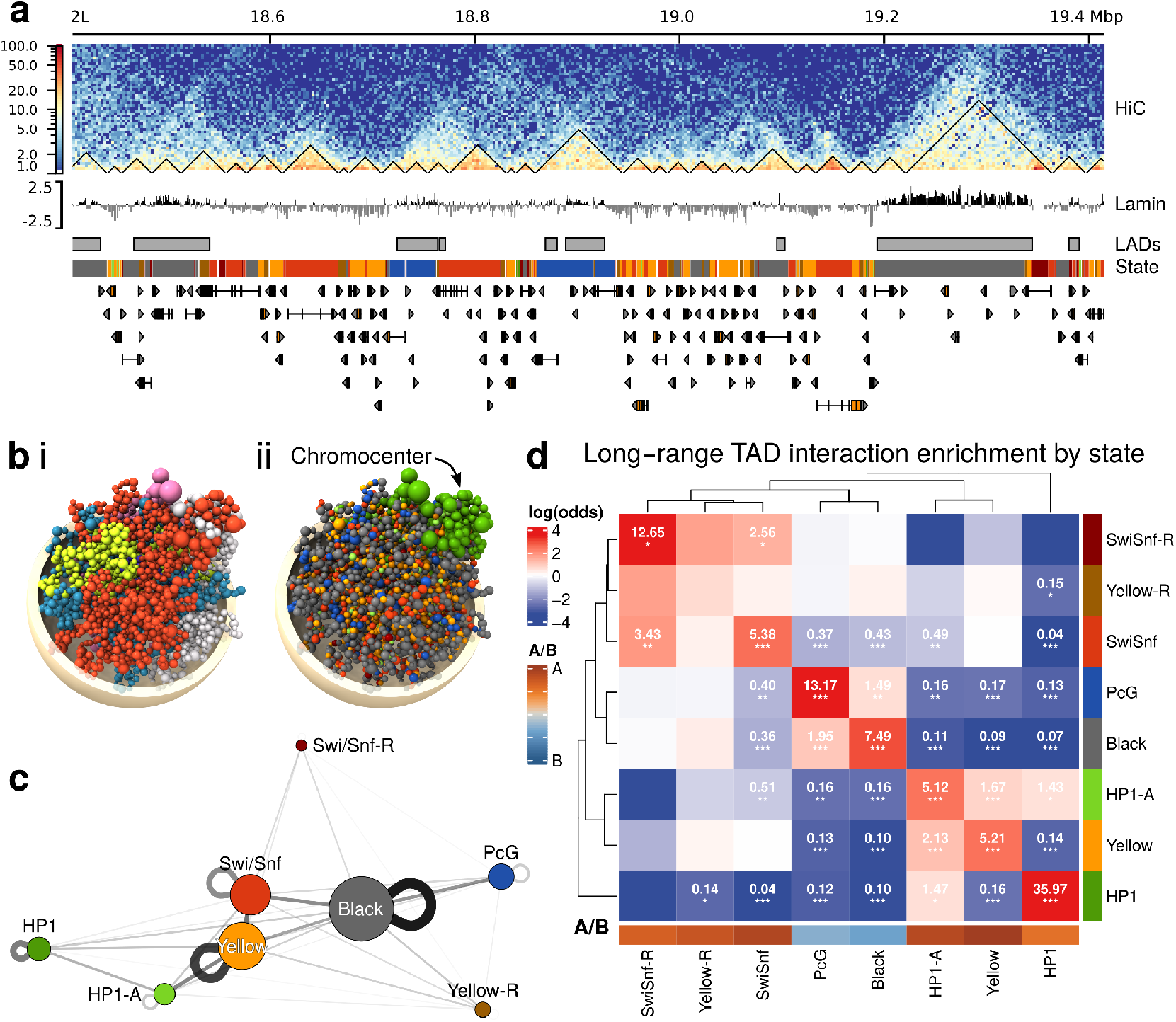
Long-range inter-TAD interactions between chromatin states compartmentalise the 3D genome. A region of the genome showing re-processed kc167 cell Hi-C data at 1kb resolution, TAD calls, Lamin-B data, LADs, chromatin states. (b) A Chrom3D model of the halploid genome, with TADs represented by spheres and coloured by (i) chromosome, and (ii) chromatin state. (c) Network graph of significantly-enriched (non-central hypergeometric distribution) inter-TAD interactions weighed by number of significant interactions; node size is proportional to the number of TADs in each state. (d) Chromatin states self-segregate and compartmentalised the 3D genome by inter-TAD interactions. Odds of association are shown for significantly enriched/depleted interactions (Fisher’s exact test P: *<0.05, **<0.01, ***<0.001, Benjamini-Hochberg adjusted) with >1.5-fold increased contact frequency; proportion of TADs in A/B compartments by chromatin state are illustrated.

Calling A/B compartments on the Hi-C matrix revealed a further layer of organisation, with PcG and Black TADs found predominantly within B compartments, and the remaining chromatin states enriched within A compartments (Fig. 6d). Surprisingly, within A compartments we observed a clear separation of states by ontology, with long-range TADs interactions divided between housekeeping/metabolic (Yellow, HP1-A, HP1) and developmental (Swi/Snf, Swi/Snf-R) roles (Fig. 6d). These data fit well with previous work in mammalian cells suggesting that B compartments were separated into two sub-compartments, and A compartments into 3 sub compartments ^46^, and suggest that a similar organisation may be found outside of *Drosophila*. Inter-TAD associations by chromatin state thus add an additional layer of complexity on the classic A/B compartmentalisation of the genome.

In conclusion, we find that chromatin is organised into eight principle states across *Drosophila* cell types. Genome organisation is divided along developmental or metabolic/-housekeeping roles, and this fundamental separation is seen across gene bodies, promoters and enhancers. Transcription factor motifs within the genome are differentially enriched based on chromatin states, and the higher-order folding of the genome is built upon chromatin state self-association, strongly suggestive of an underlying phase-separation of chromatin states.

## Methods

### Previously-published data

Details of all previously published datasets used in this study are provided in Table S1. Data were reprocessed as described below.

### Vectors, germline transformation and fly lines

pTaDaG2^53^ constructs for Brm, H1, HP1a, Su(var)3-9, Tip60 and Lam were generated by PCR amplifying the coding sequence of each chromatin factor from an embryonic cDNA library ^54^ with Gibson Assembly adaptor sequences and ligating using an NEB Hifi assembly kit (NEB). Cloning primers were designed using Perlprimer ^55^. pTaDaG2-RpII18 was generated using a synthetic gBlocks oligo (IDT) insert and NEB Hifi assembly. For profiling E(bx) and Crp binding in NSCs, we took advantage of the recently-described Nanodam technique ^28^, generating a membrane-labelled-mCherry version of the Nanodam system, *pTaDaM-traNSL-vhh4GFP*, by inserting a synthetic *traNLS-vhh4GFP* gBlock oligo (IDT) into *pTaDaM* via NEB Hifi assembly ^53^. ChromaTaDa constructs (pTaDaG2-Taf3-eCR and pTaDaG2-HP1a-eCR) were generated via the insertion of a gBlock (IDT) containing the eCR cassette into pTaDaG2 via NEB Hifi Assembly. All vectors were sequenced before being used in germline transformation.

TaDaG2-Dam, TaDaG2-Brm, TaDaG2-H1, TaDaG2-Lam, TaDaG2-RpII18, TaDaG2-Su(var)3-9, TaDaG2-Tip60 and TaDaG2-RpII18, ChromaTaDa-Taf3-eCR, ChromaTaDa-HP1a-eCR and TaDaM-traNLS-vhh4GFP fly lines were generated either via BestGene, Inc (CA), through phiC31-integrase-mediated insertion of the appropriate expression vectors into *attP2* on chromosome 3L, or via injection of *y*^*1*^*sc*^*1*^*v*^*1*^*P{nos-phiC31\int*.*NLS}X; +; P{CaryP}attP2* embryos as previously described ^8^.

Existing DamID lines used were TaDaG2-Dam ^53^, TaDaG2-Pc ^53^, and TaDaG2-MRG15 ^56^. New TaDa profiling lines for Ash2, Trr, Su(z)12 and Psc were generated as FlyORF-TaDa lines via conversion from FlyORF lines as previously described ^56^.

Gal4 driver lines were *wor-GAL4* ^57^ for neural stem cells and *R57C10-GAL4* ^58^ for mature neurons. All driver lines were crossed with *tub-Gal80ts* on either chromosome 2 or 3 to generate *w;wor-Gal4;tub-GAL80ts* and *w;tub-Gal80ts;GMR57C10-GAL4* lines respectively for use in Targeted DamID crosses. All drivers were tested for correct cell-type specific expression via crossing to the base TaDaG2-Dam line and observing the myrGFP expression pattern via confocal microscopy under the same conditions as the Targeted DamID sample collections (detailed below).

### Targeted DamID

Targeted DamID on 3rd instar larval neural stem cells and adult mature neurons was performed as previously described ^8,59,60^, using *wor-GAL4* for NSCs and *R57C10-GAL4* (commonly known as *nSyb-GAL4*) for adult neurons. Briefly, for NSCs, flies were allowed to lay on apple juice plates with yeast for 4 hours at 25°C, before transferring plates to 18°C for two days. 100 larvae from each plate were transferred to food plates and grown at 18°C for a further five days, before shifting to the permissive temperature of 29°C. Thirty brains per replicate were dissected.

For the mature neuronal dataset, flies were allowed to lay at 18°C for 24 hours in food vials. Vials were then kept at 18°C until eclosion, whereupon newly eclosed adults were transferred to fresh food vials and grown for a further day at 18°C. The vials were then shifted to 29°C for 24 hours. Following induction, flies were frozen in 14mL tubes cooled in dry ice. Frozen flies were vortexed and body parts sieved to separate the heads using mesh sizes of 710 μm, 425 μm and 150 μm; 50 heads per replicate were collected.

Following sample collection, genomic DNA was extracted, cut with DpnI and cut fragments isolated. DamID adaptors were ligated to the isolated DNA, fragments were digested with DpnII, and amplified via PCR before next-generation sequencing library preparation.

### Next-generation sequencing and data processing

Following the DamID procedure, samples were prepared for NGS as previously published ^59^. Briefly, samples were sonicated in a Bioruptor Plus (Diagenode) to reduce the average DNA fragment size to 300bp, and DamID adaptors were removed via overnight AlwI digestion. The resulting DNA was purified via magnetic bead cleanup and 500ng of DNA was end-repaired, A-tailed, ligated to NGS adaptors and amplified via 6 PCR cycles. Resultant libraries, multiplexed to yield ∼20 million mappable fragments per sample, were sequenced as either single-end reads (NextSeq500, Illumina) or paired-end reads (MGI platform, BGI).

NGS reads in FASTQ format were aligned using damidseq_pipeline ^61^ using default options for all datasets except for Histone H1 and Lamin datasets. For these, samples were normalised using RPM (reads per million) values owing to the low correlation between Dam and Dam-fusion binding profiles. The resulting gatc.bedgraph dataset replicates were compared via Pearson correlation, scaled by dividing each dataset by its standard deviation, and replicates averaged.

Gene calls from RNA Polymerase DamID-seq data were performed on averaged datasets via polii.gene.call ^59^. Datasets were visualised using pyGenomeTracks ^47^.

### ChIP-seq data reprocessing

ChIP-seq data was obtained from public repositories in FASTQ format, and aligned to release 6 of the fly genome via damidseq_pipeline (using the --chipseq parameter, which removes PCR duplicates and non-uniquely-mapping fragments), binned at either 200bp resolution (S2 cells) or GATC-fragment resolution (kc167 cells).

All binding data was transformed to a log2 binding enrichment ratio in order to fit gaussian HMMs to ChIP-seq data. For datasets with input controls, the input control reads were normalised to the binding data by the average sample / input ratio per bin, with bins in the highest two deciles of sample signal excluded. The final binding ratio for each bin was log2(sample/normalised_input). For datasets without input controls, bins with no reads were excluded and then a noise floor to the data was determined by finding the arg max of the kernel density function for the data. The final binding ratio was log2(sample / noise_floor). In both cases, pseudocounts were added per bin during ratio calculation.

### Conversion of genomic datasets to Dm6

Processed datasets (kc167 DamID, STARR-seq and STAP-seq) to Release 5 of the *Drosophila* genome were converted to the current release (Release 6) via the dmel_r5_to_r6 converter script from Flybase (https://github.com/FlyBase/bulkfile-scripts/tree/master/dmel_r5_to_r6), modified to accept files in bedgraph format.

### DamID data reprocessing

Processed, normalised microarray enrichment data for chromatin protein binding in kc167 cells (Table S1) was converted to release 6 of the *Drosophila* genome, converted to GATC resolution and scaled.

### Chromatin state modelling with ChroMATIC

ChroMATIC allows for the detection of variable levels of protein binding and histone mark occupancy via multivariate gaussian HMMs of chromatin state organisation. Unlike previous applications of HMMs to chromatin states, which have inferred states from a single best-fitted model, ChroMATIC applies multi-model inference and model averaging via relative Aikake Information Constant (AIC) weights ^62^ to a large number of full fitted models. HMMs have non-random, directional transitions between states, allowing the simple and effective visualisation of relationships between chromatin states as a weighted network graph. States are clustered by both the weighted shortest network path and by mean state emissions to assign broad chromatin groups to states for each model, together with relative probabilistic weightings, from a set of curated means. The final assigned chromatin states generated from these models represents a weighted-average fit from all models, increasing the reproducibility of HMM modelling and also providing an indication of model consistency. The ChroMATIC analysis pipeline is summarised below.

ChroMATIC takes binned protein or histone mark genomic data, expressed as a normalised log2(signal/input) ratio in bedgraph format (see above). Data are merged, sorted by chromosome and location, and gaussian Hidden Markov Models resolved via the RHmm R package (https://r-forge.r-project.org/projects/rhmm/). Given the processing demands of HMMs, models are trained on a subset of the full genomic data. For this study, either Chr2L or the full chromosome 2 (Chr2L concatenated with Chr2R) was used as the training set; this parameter is flexible, and training data can be specified as multiple chromosomes or subsets of chromosomes. Although not used in this study, when working with related datasets models can be trained on a concatenation of these data, and then applied to each dataset separately.

HMM fitting is a two-stage process. In the first stage, initialisation parameters are chosen stochastically from the range of each variable, and a large number of modelling runs are run for a set number of iterations in order to cover a broad parameter search space. We recommend 40 initial iterations as a compromise between final model accuracy and computational time. The final log likelihood (LLH) of these runs are saved, and the best-fitting parameters used to fit a full model (considered to be fitted when the change in LLH between iterations falls below a tolerance threshold). ChroMATIC is designed for use on high-performance computing (HPC) nodes with large numbers of cores. The total number of initial modelling runs is divided by the number of cores available, and each core fits a full model from its subset of parameters. We routinely use 28 cores for model fitting (thus generating 28 full models for subsequent multi-model averaging) and between 5000 and 20,000 initial modelling runs depending on dataset complexity.

In the second stage, each full model *i* is assigned an Aikake weight derived from the model Aikake Information Criterion (AIC) value ^62^ by calculating the model relative likelihood *e*^−Δ*i*/*k*^ where Δ*i* = AIC_*i*_ AIC_min_, and *k* represents a constant, and then normalising all likelihoods to sum to 1. Highly unlikely models with an Aikake weight < 0.01 are excluded from further analysis. For the remaining models, a Viterbi path is generated for each model. Fitted HMM states are matched to principal chromatin states by the correlation of state emissions to a set of curated states, and the Viterbi path reduced to principal chromatin states. A final Viterbi path is generated from the weighted average of all model Viterbi paths, using the weighted modal state at each bin. Genes are assigned chromatin states based on modal state occupancy, with the proportion of the modal state covering the gene body recorded. A high-confidence annotation of chromatin state is considered with >0.55 proportional coverage by a single state.

In addition to the final, weighted state call, ChroMATIC generates a number of additional output files representing the models. Chromatin state heatmap plots are generated via the ComplexHeatmap R package ^63^. In these plots, HMM states are clustered via kmeans and hierarchical clustering using state transition probabilities. Transcription factor coverage is based on the annotated list provided by ^37^; state expression via significantly enriched (FDR<0.01) RNA polymerase occupancy (either from DamID or ChIP-seq data) as previously described ^8,54^. Chromatin state transition network graphs were generated via the qgraph R package ^64^. For all models, a weighted graph was built from transition probabilities, and displayed via the Fruchterman-Reingold force-directed algorithm. HMM models illustrated in figures show the best-fitted model (determined via AIC) as representative; AIC-weighted average data was used in all downstream analyses.

PCA analysis of combined model data was performed on combined shared emission means and network distances of the best-fitting models. For shared emission means, binding profiles from each of the chromatin complexes listed in Table S2 (i.e. RNA Pol, Swi/Snf, H3K36me3/MRG15, HP1, PcG, and histone H1) for each cell type were isolated; the lack of H3K36me3/MRG15 profiles from ISCs meant that this cell type was excluded from this analysis. For transition network distances, fiduciary HMM states were identified for the Swi/Snf, H3K36me3/MRG15/HP1/PcG principal chromatin forms (assigned as the state in each grouping with the strongest mean emission of the relevant marker). The weighted shortest network path from each HMM state to each fiduciary state was determined via the qgraph package. Emission means and shortest paths were scaled, merged from each cell type, and PCA performed on this dataset via the prcomp R function and visualised with the ggbiplot R package. PCA variable graphs were generated using the factoextra R package.

Chromatin state transition Sankey graphs were generated using the NetworkD3 package in R.

ChroMATIC will be made available upon publication at https://github.com/marshall-lab/chromatic.

### RNA-seq data reprocessing

RNA-seq data was obtained from public repositories (Table S1) in FASTQ format and processed via Kallisto ^65^ against release 6 of the *Drosophila* genome using combined replicates; S2 cell data were paired-end and processed with default parameters; kc167 cell data were from single-end reads processed with -l 120 -s 20 parameters. TPM values per gene represent the sum of the TPM values for all transcripts of that gene.

For genome track plotting, a pseudobam coverage file was generated via Kallisto and coverage tracks at 15bp resolution generated via a custom bam2coverage script (available upon request).

### Hi-C data reprocessing

High-resolution DpnII Hi-C reads from kc167 cells (Table S1) were processed with the Hi-C explorer package ^66^. Briefly, read pairs were concatenated by paired-end to give two FASTQ files, one for each read end, and then each separately aligned to release 6 of the *Drosophila* genome, read IDs adjusted and BAM files generated for each file `$i via bowtie2 -x /mnt/genomes/dmel_release/DmR6/DmBDGP6 --threads 16 -U $i --reorder | perl -p -e ‘s/(SRR\d+\.\d+)\.\d/$1/’ | samtools view -Shb - > (basename $i .fastq.gz).bam`.

A Hi-C contact matrix was then built via hicBuildMatrix using GATC as the restriction enzyme and dangling-end sequence, and a bin size of 1kb. A matrix at 100kb resolution was also generated using hicMergeMatrixBins. After inspection of the diagnostic plots, the 1kb and 100kb matrices were separately corrected via hicCorrectMatrix.

A/B compartments were called on the corrected 1kb matrix using hicPCA. TADs were deter-mined from the 1kb matrix via hicFindTADs with the `--correctForMultipleTesting fdr parameter`.

For downstream processing with Chrom3D, matrices were converted to COOL format via hicConvertFormat.

### Hi-C A/B compartment mapping

The uncorrected Hi-C matrix was converted to 10kb resolution via hicMergeBins, and corrected. The corrected 10kb matrix was transformed to a pearson correlated matrix of log(observed/expected) interactions, and the first principle component from PCA isolated via hicPCA using gene density to determine the eigenvector sign. A/B components were assigned on positive/negative mean eigenvector values.

A/B TAD calls were merged with chromatin state calls, and all TADs with a modal state occupancy > 0.35 used for downstream analysis. For TADs of each chromatin state, the proportion of TADs in compartment A was calculated, and this value used in plotting the A/B association of each state.

### Chrom3D modelling

Chrom3D modelling was implemented as previously described ^49^ with the following modifications. Briefly, contact matrices were transformed into text matrices via cooler dump --join. Custom Rscripts were used to determine intrachromosomal contacts from the 1kb corrected matrix, and interchromosomal contacts from the 100kb corrected matrix. A custom Perl script was used to generate arrowhead domains from TADs. Enriched intrachromosomal and interchromosomal contacts, with a minimum genomic distance of 50kb, were separately determined via NCHG and NCHG_fdr_oddratio_calc.py. LADs were determined via a 2-state HMM generated with hmm.peak.caller ^56^ on existing kc167 cell Lamin-B data.

The gTrack input files for Chrom3D modeling were generated via the make_gtrack.sh script on intrachromosomal TAD contacts and arrowhead domains, then with the make_gtrack_incl_lad.sh script to add LADs, and finally the add_inter_chrom_beads.sh script to add interchromosomal contacts.

Chrom3D models then generated via multiple parallel runs of `Chrom3D -y 0.15 -r 2.0 -n 2000000 -l 10000`.

To colour TADs by chromatin state, the modal state occupancy per TAD was called via the gene.exp.modal Rscript with TAD domains passed as the gene file parameter. Chrom3D models were then recoloured by chromatin state with a custom Perl script (available upon request).

Final Chrom3D models were visualised using UCSF ChimeraX ^67^.

### Intrachromosomal long-range TAD interactions

Significantly-enriched intrachromosomal TAD interactions from NCHG (see “Chrom3D modeling”, above) were combined with chromatin state TAD calls. All interactions were filtered for log2(observed/expected)>0.58 and a modal chromatin state coverage >0.35. An enriched contact matrix of interactions between TADs of different chromatin states was then built, and enrichment of state-state interactions obtained via Fisher’s exact test. The resulting matrix of enriched state-state interactions was clustered via hclust on log2(odds of enrichment), and plotted with ComplexHeatmap ^63^.

To generate a network graph of long-range TAD interactions via chromatin state, a state-state enriched interaction matrix was generated from all interactions, weighted by cumulative log2(observed/expected) scores. The weighted graph was plotted via qgraph ^64^ using the Fruchterman-Reingold force-directed algorithm.

### Clustering of chromatin states across cell-types

Clustering was performed on all genes with a modal chromatin state occupancy >0.55 for all cell types. Each chromatin state was assigned an integer number 1:9, and clustering was performed via *k*-means with 10000 starts and a maximum of 10000 iterations per start. For ordering of clusters in the final plot, clusters were ranked by the geometric mean of chromatin states within the cluster. GO term enrichment terms were calculated per cluster via ClusterProfiler ^68^. Data on broad/specific gene expression in *Drosophila* from modEncode ^4^ was processed such that all genes with a broadly-expressed score were categorised as “broad”, and TFs ^37^ were annotated. The final plot was generated with ComplexHeatmap, using the anno_word_cloud function to generate GO term word clouds.

### Transcription factor motif enrichment analysis

Transcription factor motif enrichment was performed using the RcisTarget R package ^29^ together with the dm6-5kb-upstream-full-tx-11species.mc8nr.feather motif rankings. For each cell type studied, genes were divided by chromatin state and the cisTarget function was used to identify enriched motifs 5kb upstream of genes of the same chromatin state. For each enriched motif, the number of genes highly ranked for the motif were recorded, the coverage of the motif ([number of highly ranked genes]/[size of gene set]) within the gene set calculated, and a matrix built representing the coverage of each enriched motif per state per cell type. As a very large number of enriched motifs lack a high-confidence annotation to a TF, only motifs with a direct TF annotation were retained for plotting.

Clustering of motifs was performed with ComplexHeatmap, using Spearman’s rank correlation as the distance method between rows and columns, clustering rows and columns via centroid heirarchical clustering and splitting columns on chromatin state; given the very large number of specifically-enriched motifs at low coverage, the matrix was filtered for a minimum motif enrichment coverage of 0.05 prior to plotting.

## STARR-seq data analysis

Processed S2 STARR-seq merged-peaks data for both dCPs and hkCPs (Table S1) were converted to Dm6 genome coordinates, putative enhancers in each set with an activation score < 5 were considered inactive and filtered, and the chromatin state coverage around each active enhancer center +/-50bp was determined, together with the average occupancy of all chromatin proteins and histone marks. Enrichment of chromatin state coverage per enhancer type was calculated via Fisher’s exact test.

## STAP-seq data analysis

Processed STAP-seq data for core promoters (Table S1) were converted to Dm6 genome coordinates and the binding data z-transformed via scaling the translated matrix (such that scaling was performed on the transcriptional co-factor (CoF) binding data for each promoter). As the published clustering of core promoters was not available, clustering of the complete CoF dataset was performed via *k*-means for 5 clusters, using 1000 starts, a maximum of 1000 iterations per start, and the MacQueen *k*-means algorithm (due to Quick-TRANSfer convergence failure of the standard Hartigan-Wong algorithm). As the published clusters showed specific enrichment for p65, Med25, Mof, Chro and gfzf, the median CoF binding per cluster for these factors was compared per cluster, and cluster numbers assigned such that they accorded to the published clustering numbers. Each core promoter was assigned a chromatin state based on the modal occupancy of the core promoter center +/-50bp. Enrichment of chromatin state occupancy was determined via Fisher’s exact test.

Processed STAP-seq data for specific enhancer activation per core promoter (using the *zfh1* enhancer for developmental enhancer activation, and the *ssp3* enhancer for house-keeping activation) were converted to Dm6 coordinates. The signal for each enhancer was z-transformed by scaling, and data were filtered for core promoters that were specifically activated by only one enhancer, such that their activation z-score was > 0 for one enhancer type and < 0 for the other. These specifically-activated core-promoters were then plotted by chromosome state per cluster.

### Larval brain scRNA-sequencing reanalysis

Seurat v4^69^ was used to manually annotate the Seurat object from a published scRNA-seq dataset of the larval brain ^39^, and identify NSCs, cholinergic, glutamatergic, and GABAergic neuronal clusters using the established markers indicated in Fig. 5d. The DotPlot function from Seurat was used to extract percent and scaled average gene expression values for the selected clusters. Genes were filtered by chromatin state and ranked by FDR and fold change based on RpII18 (RNA Polymerase) occupancy, and the final plot generated with ComplexHeatmap ^63^.

### Genomic features

Genomic features (exons, introns and intergenic regions) were obtained via the following method. The *D. melanogaster* release 6.34 GTF annotations were read as a GRanges object, filtered for exons, and set as unstranded. The exon Granges object was reduced, and all non-exon regions taken as the gaps in this object. Separately, the R genomation package was used to read features from the same genome GTF file, identifying introns. Intergenic regions were taken to be the non-exon ranges with the intron ranges excluded.

### Statistical tests

All P-values were Benjamini-Hochberg corrected, and alpha was assigned as 0.05 unless otherwise specified.

### Gene ontogeny analysis

All gene ontogeny (GO) term analysis was carried out via the ClusterProfiler R package ^68^.

### Other bioinformatics analyses

Bioinformatics pipelines were accelerated using GNU Parallel ^70^. All other analyses were performed using R ^71^.

## Software availability

All analysis code used in this project will be made available on GitHub at https://github.com/marshall-lab upon publication.

## Data availability

Next-generation sequencing data generated in this study will be deposited in NCBI GEO prior to publication.

## Acknowledgments

We thank Grace Jefferies, Victoria Roy and Elizabeth Read for technical assistance. We thank Ciarán O’Mara, Jake Newland and all members of the Marshall Group, past and present, for their helpful thoughts, comments, insights and discussions. We also thank Alison Bardin, Natalia Rubanova, Julie Secombe and Angela Giangrande for interesting and stimulating discussions on the nature of chromatin. This work was supported by NHMRC grants APP1128784 and APP185220, Ian Potter Foundation grant 20190091 and philanthropic funding from the Menzies Institute for Medical Research, to OJM.

## Author contributions

OJM conceived and designed the research; CD, JPDM and OJM performed experiments and obtained the data; OJM wrote the software; OJM, CD, PC and JP analysed the data; OJM, CD and PC wrote the manuscript.

## References

1. Ernst J and Kellis M. Nature biotechnology, 28(8):817–25, 2010.

2. Kharchenko PV, Alekseyenko AA, Schwartz YB, et al. Nature, 471(7339):480–485, 2011.

3. Ernst J, Kheradpour P, Mikkelsen TS, et al. Nature, 473(7345):43–9, 2011.

4. Ho JWK, Jung YL, Liu T, et al. Nature, 512(7515):449–52, 2014.

5. Consortium RE, Kundaje A, Meuleman W, et al. Nature, 518(7539):317–330, 2015.

6. Filion GJ, van Bemmel JG, Braunschweig U, et al. Cell, 143(2):212–24, 2010.

7. van Bemmel JG, Filion GJ, Rosado A, et al. Molecular Cell, 49(4):759–771, 2013.

8. Marshall OJ and Brand AH. Nature Communications, 8(1):2271, 2017.

9. Delandre C and Marshall OJ. Biochemical Society Transactions, page BST20180605, 2019.

10. Shao Z, Raible F, Mollaaghababa R, et al. Cell, 98(1):37–46, 1999.

11. Marenda DR, Zraly CB, and Dingwall AK. Developmental Biology, 267(2):279–293, 2004.

12. Lavigne M, Eskeland R, Azebi S, et al. PLoS Genet, 5(12):e1000769, 2009.

13. Hahn MA, Wu X, Li AX, et al. PLoS ONE, 6(4):e18844, 2011.

14. Riddle NC, Minoda A, Kharchenko PV, et al. Genome Res., 21(2):147–163, 2011.

15. Schmitges FW, Prusty AB, Faty M, et al. Molecular Cell, 42(3):330–341, 2011.

16. Mauser R, Kungulovski G, Keup C, et al. Epigenetics & Chromatin, 10(1):45, 2017.

17. Barral A, Pozo G, Ducrot L, et al. Molecular Cell, 82(4):816–832.e12, 2022.

18. Harikrishnan KN, Chow MZ, Baker EK, et al. Nature Genetics, 37(3):254–264, 2005.

19. Zhang X, Li B, Li W, et al. Stem Cell Reports, 3(3):460–474, 2014.

20. Raab JR, Resnick S, and Magnuson T. PLoS Genetics, 11(12):1–26, 2015.

21. Riddle NC, Jung YL, Gu T, et al. PLoS genetics, 8(9):e1002954, 2012.

22. Becker JS, McCarthy RL, Sidoli S, et al. Molecular Cell, 68(6):1023–1037.e15, 2017.

23. Kolasinska-Zwierz P, Down T, Latorre I, et al. Nature Genetics, 41(3):376–381, 2009.

24. Arnold CD, Gerlach D, Stelzer C, et al. Science (New York, N.Y.), 339(6123):1074–7, 2013.

25. Haberle V, Arnold CD, Pagani M, et al. Nature, 4, 2019.

26. Hendy O, Serebreni L, Bergauer K, et al. Molecular Cell, page S1097276522008085, 2022.

27. Fazzio TG, Huff JT, and Panning B. Cell, 134(1):162–174, 2008.

28. Tang JL, Hakes AE, Krautz R, et al. Developmental Cell, 57:1–15, 2022.

29. Davie K, Janssens J, Koldere D, et al. Cell, 174(4):982–998.e20, 2018.

30. Wong MMK, Liu MF, and Chiu SK. BMC Dev Biol, 15(1):20, 2015.

31. Yang J, Ramos E, and Corces VG. Genome Res., 22(11):2199–2207, 2012.

32. Chang KC, Garcia-Alvarez G, Somers G, et al. Developmental Cell, 19(5):778–785, 2010.

33. Touma JJ, Weckerle FF, and Cleary MD. Development (Cambridge, England), 139(4):657–66, 2012.

34. Dalton D, Chadwick R, and McGinnis W. Genes & Development, 3(12a):1940–1956, 1989.

35. Wang L and Coulter DE. The EMBO Journal, 15(12):3182–3196, 1996.

36. Dallman JE, Allopenna J, Bassett A, et al. The Journal of neuroscience : the official journal of the Society for Neuroscience, 24(32):7186–7193, 2004.

37. Hens K, Feuz JD, Isakova A, et al. Nature methods, 8(12):1065–70, 2011.

38. Schwartz YB, Kahn TG, Nix DA, et al. Nature Genetics, 38(6):700–705, 2006.

39. Dillon N, Cocanougher B, Sood C, et al. Neural Dev, 17(1):7, 2022.

40. Featherstone DE, Rushton E, and Broadie K. Nat Neurosci, 5(2):141–146, 2002.

41. Kern JV, Zhang YV, Kramer S, et al. Genetics, 195(1):59–72, 2013.

42. Han M, Park D, Vanderzalm PJ, et al. Journal of Neurochemistry, 90(1):129–141, 2004.

43. Mezler M, Müller T, and Raming K. European Journal of Neuroscience, 13(3):477–486, 2001.

44. Szabo Q, Jost D, Chang JM, et al. Science advances, 4(2):eaar8082, 2018.

45. Lieberman-Aiden E, van Berkum NL, Williams L, et al. Science (New York, N.Y.), 326(5950):289–93, 2009.

46. Rao SS, Huntley MH, Durand NC, et al. Cell, 159(7):1665–1680, 2014.

47. Cubeñas-Potts C, Rowley MJ, Lyu X, et al. Nucleic acids research, 45(4):1714–1730, 2017.

48. Paulsen J, Sekelja M, Oldenburg AR, et al. Genome Biology, 18(1):1–15, 2017.

49. Paulsen J, Liyakat Ali TM, and Collas P. Nature Protocols, 13(5):1137–1152, 2018.

50. Strom AR, Emelyanov AV, Mir M, et al. Nature, 547(7662):241–245, 2017.

51. Gibson BA, Doolittle LK, Schneider MWG, et al. Cell, pages 1–15, 2019.

52. Seif E, Kang JJ, Sasseville C, et al. Nature Communications, 11(1):1–19, 2020.

53. Delandre C, McMullen JPD, and Marshall OJ. bioRxiv, page doi:10.1101/2020.04.17.045948.

54. Southall TD, Gold KS, Egger B, et al. Developmental cell, 26(1):101–12, 2013.

55. Marshall OJ. Bioinformatics, 20(15):2471–2, 2004.

56. Aughey GN, Delandre C, McMullen JPD, et al. G3 Genes|Genomes|Genetics, 11(1):1–6, 2021.

57. Albertson R, Chabu C, Sheehan A, et al. Journal of cell science, 117(Pt 25):6061–70, 2004.

58. Henry GL, Davis FP, Picard S, et al. Nucleic Acids Research, 40(19):9691–9704, 2012.

59. Marshall OJ, Southall TD, Cheetham SW, et al. Nature Protocols, 11(9):1586–1598, 2016.

60. Marshall OJ and Delandre C. In MJ Horsfield J, editor, Chromatin. Methods in Molecular Biology, volume 2458, pages 195–213. Humana, New York, NY, 2022. ISBN 978-1-07-162139-4.

61. Marshall OJ and Brand AH. Bioinformatics (Oxford, England), 31(20):3371–3, 2015.

62. Burnham KP and Anderson DR. Sociological Methods and Research, 33(2):261–304, 2004.

63. Gu Z, Eils R, and Schlesner M. Bioinformatics, 32(18):2847–2849, 2016.

64. Epskamp S, Cramer AOJ, Waldorp LJ, et al. J. Stat. Soft., 48(4), 2012.

65. Bray NL, Pimentel H, Melsted P, et al. Nat Biotechnol, 34(5):525–527, 2016.

66. Ramírez F, Bhardwaj V, Arrigoni L, et al. Nature Communications, 9(1), 2018.

67. Goddard TD, Huang CC, Meng EC, et al. Protein Science, 27(1):14–25, 2018.

68. Yu G, Wang LG, Han Y, et al. OMICS A Journal of Integrative Biology, 16(5):284–287, 2012.

69. Hao Y, Hao S, Andersen-Nissen E, et al. Cell, 184(13):3573–3587.e29, 2021.

70. Tange O. ;login: The USENIX Magazine, 36(1):42–47, 2011.

71. R_Development_Core_Team. R: A Language and Environment for Statistical Computing, 2011.

72. Gervais L, van den Beek M, Josserand M, et al. Developmental Cell, 49(4):556–573, 2019.

73. Jordán-Pla A, Yu S, Waldholm J, et al. BMC Genomics, 19(1):1–12, 2018.

74. Climent-Canto P, Carbonell A, Tatarski M, et al. Nucleic Acids Research, 48(8):4147–4160, 2020.

75. Henriques T, Scruggs BS, Inouye MO, et al. Genes and Development, 32(1):26–41, 2018.

76. Colmenares SU, Swenson JM, Langley SA, et al. Developmental Cell, 42(2):156–169.e5, 2017.

77. Huang C, Yang F, Zhang Z, et al. Nature Communications, 8(1), 2017.

78. Heurteau A, Perrois C, Depierre D, et al. Genome Biology, 21(1):1–19, 2020.

79. Schauer T, Ghavi-Helm Y, Sexton T, et al. EMBO reports, 18(10):1854–1868, 2017.

80. Albig C, Wang C, Dann GP, et al. Nature Communications, 10(1):1–17, 2019.

81. Alekseyenko AA, Gorchakov AA, Zee BM, et al. Genes and Development, 28(13):1445–1460, 2014.

82. Enderle D, Beisel C, Stadler MB, et al. Genome Research, 21(2):216–226, 2011.

83. Tchurikov NA, Klushevskaya ES, Fedoseeva DM, et al. Cells, 9(12), 2020.

84. Samata M, Alexiadis A, Richard G, et al. Cell, 182(1):127–144.e23, 2020.

85. Zabidi MA, Arnold CD, Schernhuber K, et al. Nature, 518(7540):556–559, 2015.

86. Kellner WA, Ramos E, Van Bortle K, et al. Genome Research, 22(6):1081–1088, 2012.

87. Li L, Lyu X, Hou C, et al. Molecular Cell, 58(2):216–231, 2015.

88. Nazer E, Dale RK, Chinen M, et al. PLoS Genet, 14(3):e1007276, 2018.

89. Rowley MJ, Nichols MH, Lyu X, et al. Molecular Cell, 67(5):837–852.e7, 2017.

90. Villa R, Jagtap PKA, Thomae AW, et al. Genes Dev., 35(13-14):1055–1070, 2021.

91. Boettiger AN, Bintu B, Moffitt JR, et al. Nature, 529(7586):418–422, 2016.

92. Akhtar J, More P, Albrecht S, et al. Life Sci. Alliance, 2(4):e201900318, 2019.

93. Villaseñor R, Pfaendler R, Ambrosi C, et al. Nature Biotechnology, 38(6):728–736, 2020.

