## Supplementary data for "Eight principal chromatin states functionally segregate the fly genome into developmental and housekeeping roles"

### Supplementary Tables

**Table S1**: Accession numbers and sources of genomic data used in this study.

| Protein / mark | Method | Data source | Reference |
|---|---|---|---|
| **Larval NSCs** | | | |
| Ash2 | FlyORF-TaDa | [TBA] | [this study] |
| Brm | TaDa | [TBA] | [this study] |
| Cropped | Nanodam | [TBA] | [this study] |
| E(bx) | Nanodam | [TBA] | [this study] |
| H1 | TaDa | [TBA] | [this study] |
| H3K4me3-eCR | ChromaTaDa | [TBA] | [this study] |
| H3K9me3-eCR | ChromaTaDa | [TBA] | [this study] |
| HP1a | TaDa | [TBA] | [this study] |
| Lamin | TaDa | [TBA] | [this study] |
| MRG15 | TaDa/FlyORF-TaDa | GSE159632 | Ref. 56 |
| Pc | TaDa | [TBA] | Ref. 53 |
| Psc | FlyORF-TaDa | [TBA] | [this study] |
| RNA Polymerase (all types) (RpII18) | TaDa | [TBA] | [this study] |
| Su(var)3-9 | TaDa | [TBA] | [this study] |
| Su(z)12 | FlyORF-TaDa | [TBA] | [this study] |
| Trr | FlyORF-TaDa | [TBA] | [this study] |
| **Adult neurons** | | | |
| Brm | TaDa | [TBA] | [this study] |
| H1 | TaDa | [TBA] | [this study] |
| HP1a | TaDa | [TBA] | [this study] |
| Lamin | TaDa | [TBA] | [this study] |
| MRG15 | TaDa | [TBA] | [this study] |
| Pc | TaDa | [TBA] | [this study] |
| RNA Polymerase (all types) (RpII18) | TaDa | [TBA] | [this study] |
| Su(var)3-9 | TaDa | [TBA] | [this study] |
| Tip60 | TaDa | [TBA] | [this study] |
| **Adult ISCs** | | | |
| Brm | TaDa | GSE128941 | Ref. 72 |
| H1 | TaDa | GSE128941 | Ref. 72 |
| HP1 | TaDa | GSE128941 | Ref. 72 |
| Pc | TaDa | GSE128941 | Ref. 72 |
| RNA Polymerase II (RpII215) | TaDa | GSE128941 | Ref. 72 |
| Trr | TaDa | GSE128941 | Ref. 72 |
| **S2 Cells** | | | |
| Brm | ChIP-seq | GSE95236 | Ref. 73 |
| H1 | ChIP-seq | GSE127224 | Ref. 74 |
| H3K4me1 | ChIP-seq | GSE85191 | Ref. 75 |
| H3K4me3 | ChIP-seq | GSE85191 | Ref. 75 |
| H3K9me3 | ChIP-seq | GSE99027 | Ref. 76 |
| H3K27ac | ChIP-seq | GSE85191 | Ref. 75 |

| Protein / mark | Method | Data source | Reference |
|---|---|---|---|
| H3K27me3 | ChIP-seq | GSE93100 | Ref. 77 |
| | ChIP-seq | GSE130211 | Ref. 78 |
| H3K36me2 | ChIP-seq | GSE93100 | Ref. 77 |
| H3K36me3 | ChIP-seq | GSE94115 | Ref. 79 |
| | ChIP-seq | GSE99027 | Ref. 76 |
| | ChIP-seq | GSE128455 | Ref. 80 |
| H4K16ac | ChIP-seq | GSE94115 | Ref. 79 |
| HP1a | ChIP-seq | GSE56101 | Ref. 81 |
| ISWI | CUT&RUN | GSE184187 | Ref. 26 |
| Mrg15 | ChIP-seq | GSE93100 | Ref. 77 |
| Pc | ChIP-seq | GSE24521 | Ref. 82 |
| RNA Polymerase II | ChIP-seq | GSE85191 | Ref. 75 |
| Snr1 | CUT&RUN | GSE184187 | Ref. 26 |
| **S2 Cells additional** | | | |
| RNA-seq | | GSE145320 | Ref. 83 |
| | | GSE130333 | Ref. 84 |
| STARR-seq | | GSE57876 | Ref. 85 |
| STAP-seq | | https://starklab.org/data/ | Ref. 25 |
| **kc167 cells** | | | |
| Brm | DamID | GSE36175 | Ref. 6,7 |
| Caf1 | DamID | GSE36175 | Ref. 6,7 |
| E(z) | DamID | GSE36175 | Ref. 6,7 |
| H1 | DamID | GSE36175 | Ref. 6,7 |
| H3K4me1 | ChIP-seq | GSE36374 | Ref. 86 |
| | ChIP-seq | GSE62904 | Ref. 87 |
| | ChIP-seq | GSE95402 | Ref. 88 |
| H3K4me3 | ChIP-seq | GSE62904 | Ref. 87 |
| | ChIP-seq | GSE95402 | Ref. 88 |
| H3K9me2 | ChIP-seq | GSE62904 | Ref. 87 |
| H3K9me3 | ChIP-seq | GSE89244 | Ref. 89 |
| H3K27ac | ChIP-seq | GSE36374 | Ref. 86 |
| | ChIP-seq | GSE95402 | Ref. 88 |
| H3K27me3 | ChIP-seq | GSE80702 | Ref. 47 |
| H3K36me3 | ChIP-seq | GSE89244 | Ref. 89 |
| | ChIP-seq | GSE128455 | Ref. 80 |
| H4K16ac | ChIP-seq | GSE89244 | Ref. 89 |
| | ChIP-seq | GSE165833 | Ref. 90 |
| H4K20me1 | ChIP-seq | GSE89244 | Ref. 89 |
| HP1a | DamID | GSE36175 | Ref. 6,7 |
| Iswi | DamID | GSE36175 | Ref. 6,7 |
| Jasper | DamID | GSE36175 | Ref. 6,7 |
| Lamin | DamID | GSE36175 | Ref. 6,7 |
| MRG15 | DamID | GSE36175 | Ref. 6,7 |
| Pc | DamID | GSE36175 | Ref. 6,7 |
| RNA Polymerase (all types) (RpII18) | DamID | GSE36175 | Ref. 6,7 |
| Sce | DamID | GSE36175 | Ref. 6,7 |

| Protein / mark | Method | Data source | Reference |
|---|---|---|---|
| Sir2 | DamID | GSE36175 | Ref. 6,7 |
| Su(var)3-9 | DamID | GSE36175 | Ref. 6,7 |
| **kc167 cells additional** | | | |
| Hi-C (DpnII) | | GSE80702 | Ref. 47 |
| RNA-seq | | GSE75060 | Ref. 91 |
| **NSC tumours** | | | |
| H3K4me3 | ChIP-seq | GSE112633 | Ref. 92 |
| **Larval brain scRNA-seq** | | | |
| atlas_combined.rds Seurat object | scRNA-seq | Data link | Ref. 39 |

**Table S2**: Protein and histone mark coverage of major chromatin complexes profiled in this study, by cell type

| **Cell type** | **RNA pol**[a] | **TrxG** | | | **H3K36me3-related** | **HP1** | **PcG** | | **Repressive** |
|---|---|---|---|---|---|---|---|---|---|
| | | Swi/Snf | NuRF | COMPASS | | | PRC1 | PRC2 | |
| NSCs | RpII18 | Brm | E(bx) | Ash2, Trr, H3K4me3 | MRG15 | HP1a, Su(var)3-9, H3K9me3 | Pc, Psc | Su(z)12, H3K27me3 | H1, Lamin |
| kc167 | RpII18 | Brm | ISWI | H3K4me3 | MRG15, Jasper, H3K36me3 | HP1a, Su(var)3-9, H3K9me3 | Pc, Sce | E(z), H3K27me3 | H1, Lamin |
| S2 | RNA pol II | Brm,Snr1 | ISWI | H3K4me3 | MRG15, Jasper, H3K36me3 | HP1a, H3K9me3 | Pc | H3K27me3 | H1 |
| Neurons | RpII18 | Brm | | | MRG15, Tip60 | HP1a | Pc | | H1 |
| ISCs | RpII215 | Brm | | Trr | | HP1a | Pc | | H1 |

[a]RNA polymerase profiled either via DamID of RpII18 (shared subunit of all three RNA polymerases), DamID of RpII215 (subunit of RNA Pol II) or ChIP-seq of RNA Pol II.

### Supplementary Figures

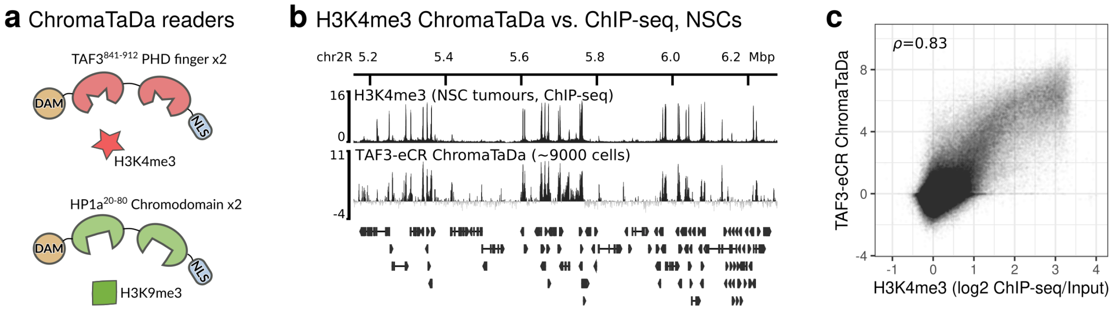

**Figure S1:** ChromaTaDa allows sensitive, cell-type specific profiling of histone modification marks *in vivo*. (a) ChromaTaDa histone mark readers were designed along the principles of the eCR readers – isolated multimers of residue-specific chromodomains or PHD fingers – from the ChromID system[93]. For profiling H3K4me3, the PHD finger (residues 841-912)[93] from mouse TAF3 was used; for profilng H3K9me3, the chromodomain from *Drosophila* HP1a (residues 20-80) was used. The final ChromaTaDa profiling proteins are formed from two identical histone-mark-reading motifs, linked by a flexible 30aa Gly-rich linker region, and fused to DNA Adenine Methyltransferase (DAM) from *E. coli* at the N-terminal and a nuclear localisation sequence (NLS) on the C-terminal. (b) ChromaTaDa profiling of H3K4me3 in NSCs compared against previously-published H3K4me3 ChIP-seq profiling in NSC-enriched larval brain tumours shows (c) high correlation of the two binding profiles (ChIP-seq profile transformed as a log2(signal/input) ratio – see Methods for details).

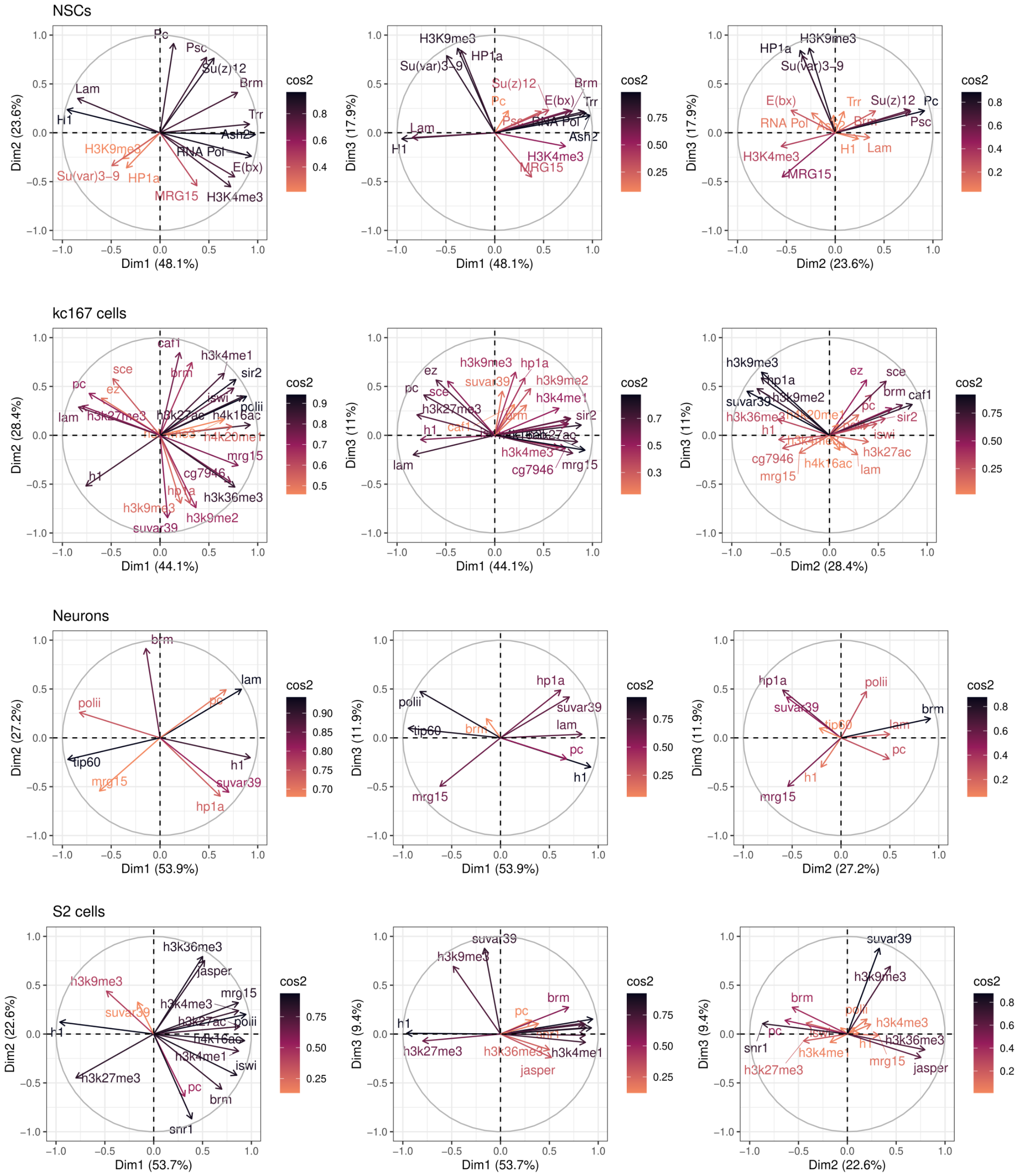

**Figure S2:** PCA variable graphs from ChroMATIC state emissions, illustrating contributions of chromatin proteins and marks to state emission means for the first three principal components of each analysis, and coloured by variable quality (cos2). Graph vectors show the contribution of each variable to each principal component dimension (Dim) through proximity to the unit circle; the angle between two vectors represents the correlation between two variables.

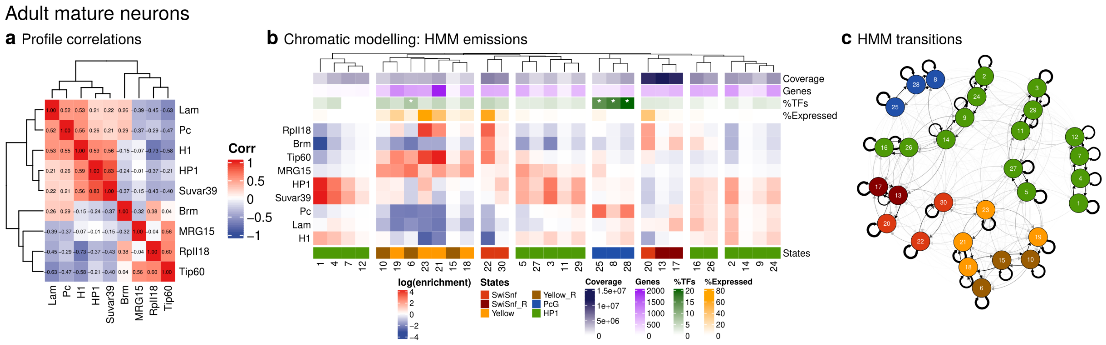

**Figure S3:** Chromatic models of chromatin states from adult mature neurons. (a) Correlation of all binding profiles used for this cell line. (b) HMM state emissions heatmap; genomic coverage, number of genes, percentage of genes as TFs per state (* = P<0.01, Fisher's exact test, Benjamini-Hochberg adjusted), and percentage of genes expressed, is also shown. (b) HMM transitions network graph. The best-fitting model based on relative AIC weighting is shown; final state assignments were derived from a weighted-average of all fitted models.

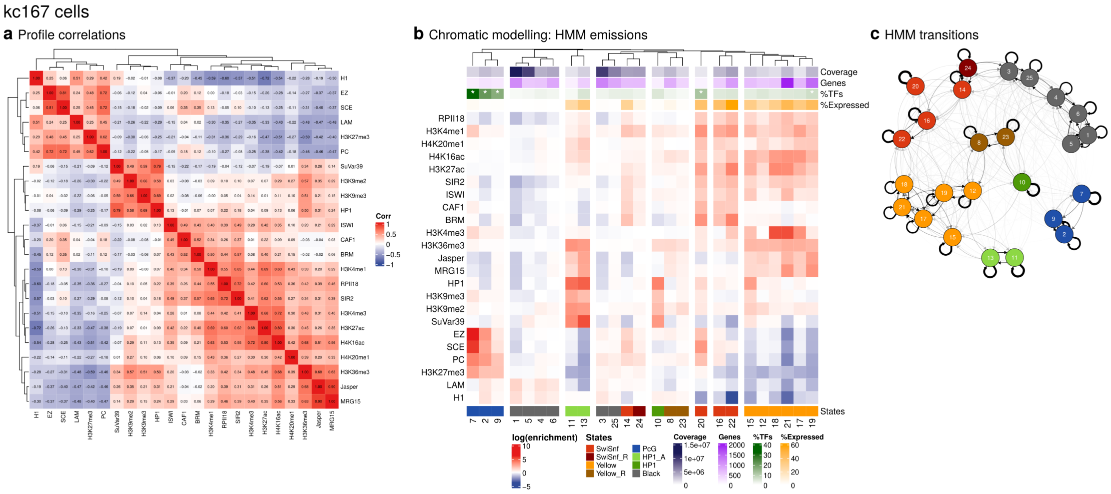

**Figure S4:** Chromatic models of chromatin states from kc167 cells. (a) Correlation of all binding profiles used for this cell line. (b) HMM state emissions heatmap; genomic coverage, number of genes, percentage of genes as TFs per state (* = P<0.01, Fisher's exact test, Benjamini-Hochberg adjusted), and percentage of genes expressed, is also shown. (c) HMM transitions network graph. The best-fitting model based on relative AIC weighting is shown; final state assignments were derived from a weighted-average of all fitted models.

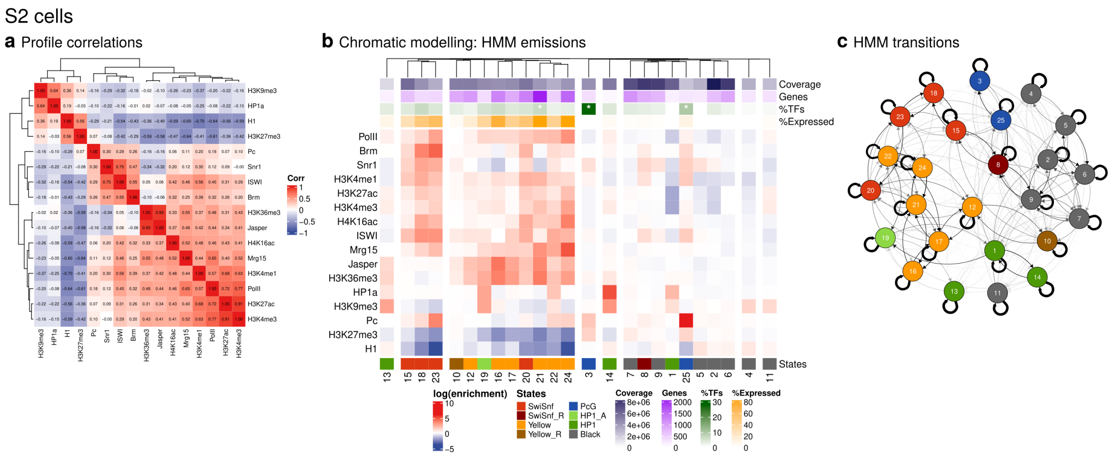

**Figure S5:** Chromatic models of chromatin states from S2 cells. (a) Correlation of all binding profiles used for this cell line. (b) HMM state emissions heatmap; genomic coverage, number of genes, percentage of genes as TFs per state (* = P<0.01, Fisher's exact test, Benjamini-Hochberg adjusted), and percentage of genes expressed, is also shown. (c) HMM transitions network graph. The best-fitting model based on relative AIC weighting is shown; final state assignments were derived from a weighted-average of all fitted models.

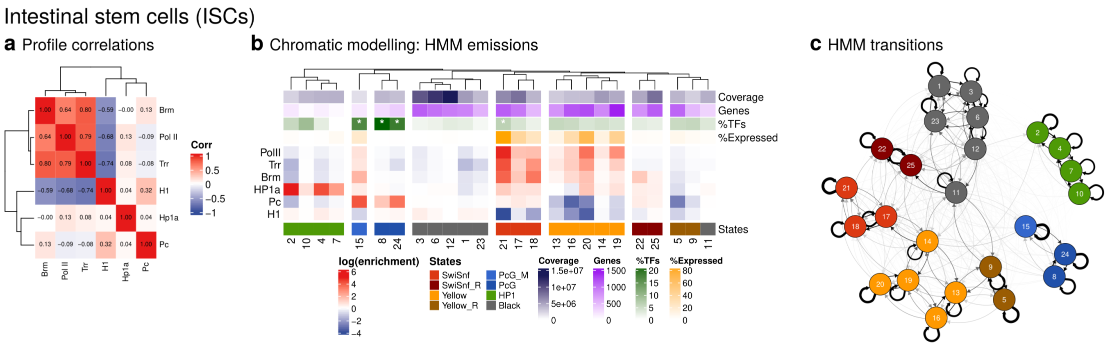

**Figure S6:** Chromatic models of chromatin states from intestinal stem cells (ISCs). (a) Correlation of all binding profiles used for this cell line. (b) HMM state emissions heatmap; genomic coverage, number of genes, percentage of genes as TFs per state (* = P<0.01, Fisher's exact test, Benjamini-Hochberg adjusted), and percentage of genes expressed, is also shown. (c) HMM transitions network graph. The best-fitting model based on relative AIC weighting is shown; final state assignments were derived from a weighted-average of all fitted models. For ISCs, no data for the binding of Yellow chromatin components was available, but we took advantage of the refractory relationship between Pc and MRG15 present in other datasets to infer Yellow/Yellow-R chromatin from Pc-depleted clusters of states.

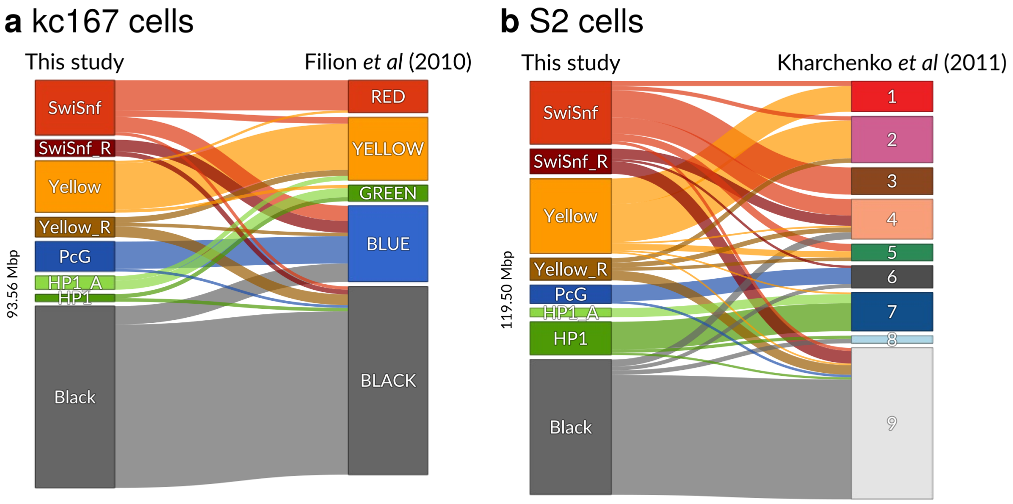

**Figure S7:** Comparison of Chromatin state modelling against previously-published chromatin state modelling for (a) kc167 cells (5 state *t*-distributed HMM of first three PCA components[6]) and (b) S2 cells (9 state gaussian HMM of 18 histone mark modifications[2]). Sankey networks of genomic coverage are shown, based on GATC-fragment level bins for kc167 cells, and 200bp binds for S2 cells. The difference in genome coverage between the two cell types reflects the relative lack of coverage for heterochromatic regions in the microarray-derived kc167 dataset. For visual simplification, transitions in the Sankey networks covering less than 500kb of the genome are hidden. State names and colours are as per the original publications.

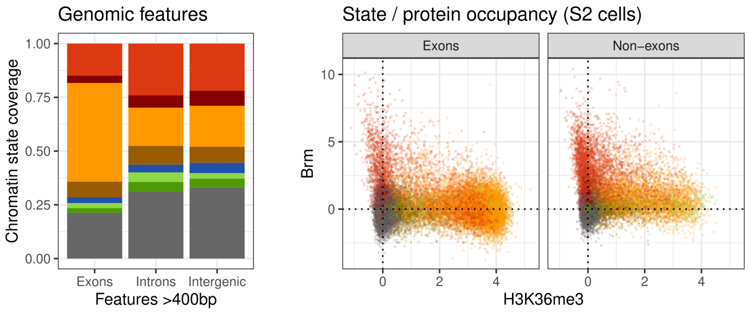

**Figure S8:** Genomic feature coverage and per-feature individual Brm/H3K36me3 occupancy values for S2 cells, showing clear separation of Brm and H3K36me3 occupancy over exonic and non-exonic regions. To exclude signal overlap, only features >400bp are shown.

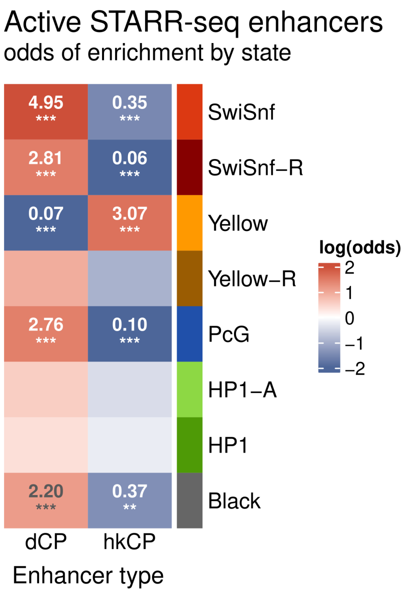

**Figure S9:** Chromatin state enrichment of active enhancers that drive either developmental (dCP) or housekeeping (hkCP) core promoters. Odds of association are shown for significantly enriched/depleted interactions (Fisher's exact test P: *<0.05, **<0.01, ***<0.001, Benjamini-Hochberg adjusted)

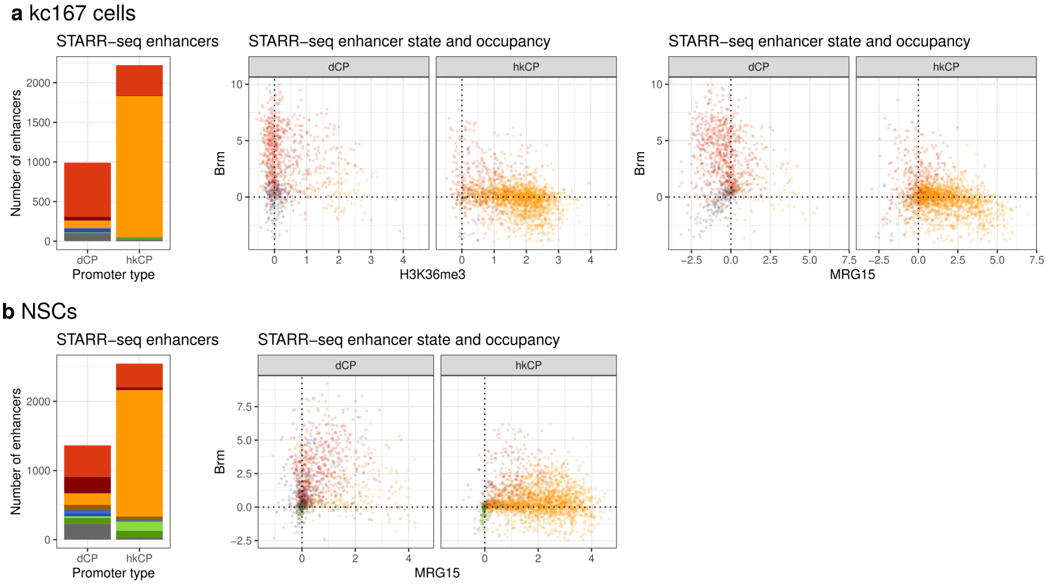

**Figure S10:** Chromatin state and individual enhancer region occupancy for developmental or housekeeping specific STARR-seq enhancers in S2 cells, using (a) kc167 chromatin states and protein/mark occupancy; and (b) NSC state and protein occupancy data. In each cell type, unrelated to S2 cells, the separation between developmental / housekeeping states, and Brm / MRG15-H3K36me3 is shown, illustrating a consistent separation between Brm and MRG15/H3K36me3 occupancy over developmental and housekeeping enhancers.

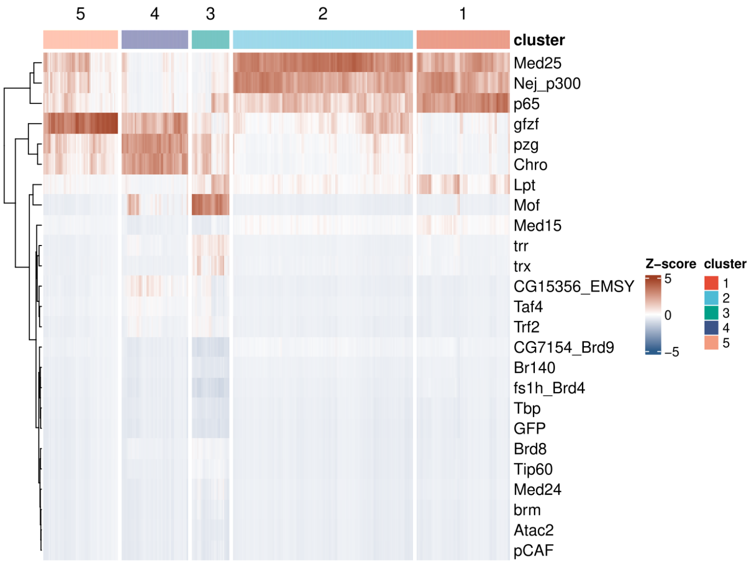

**Figure S11:** Re-analysis of STAP-seq core promoter (CP) activation data[25] to re-derive the 5 distinct CP clusters, numbered and coloured as per[25]; see Methods for re-analysis details.

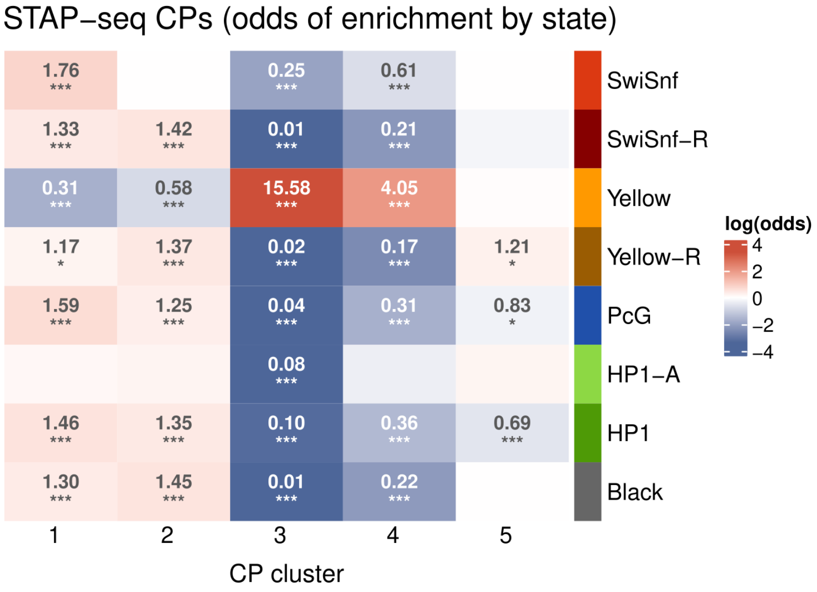

**Figure S12:** Chromatin state enrichment of different core promoter classes (clusters are annotated as per Fig. S11). Odds of association are shown for significantly enriched/depleted interactions (Fisher's exact test P: *<0.05, **<0.01, ***<0.001, Benjamini-Hochberg adjusted)

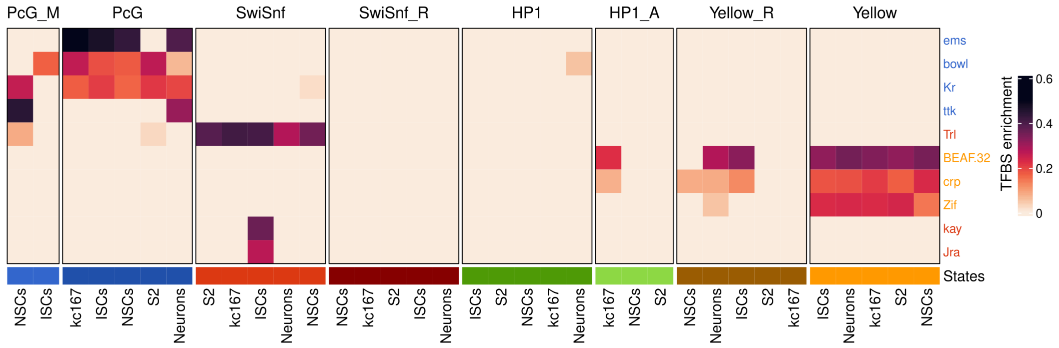

**Figure S13:** Details of transcription factor binding site (TFBS) analysis from Fig. 3; only highlighted genes are illustrated. Final binding site enrichment values represent the proportion of genes in a chromatin state with an enrichment of the TFBS motif.

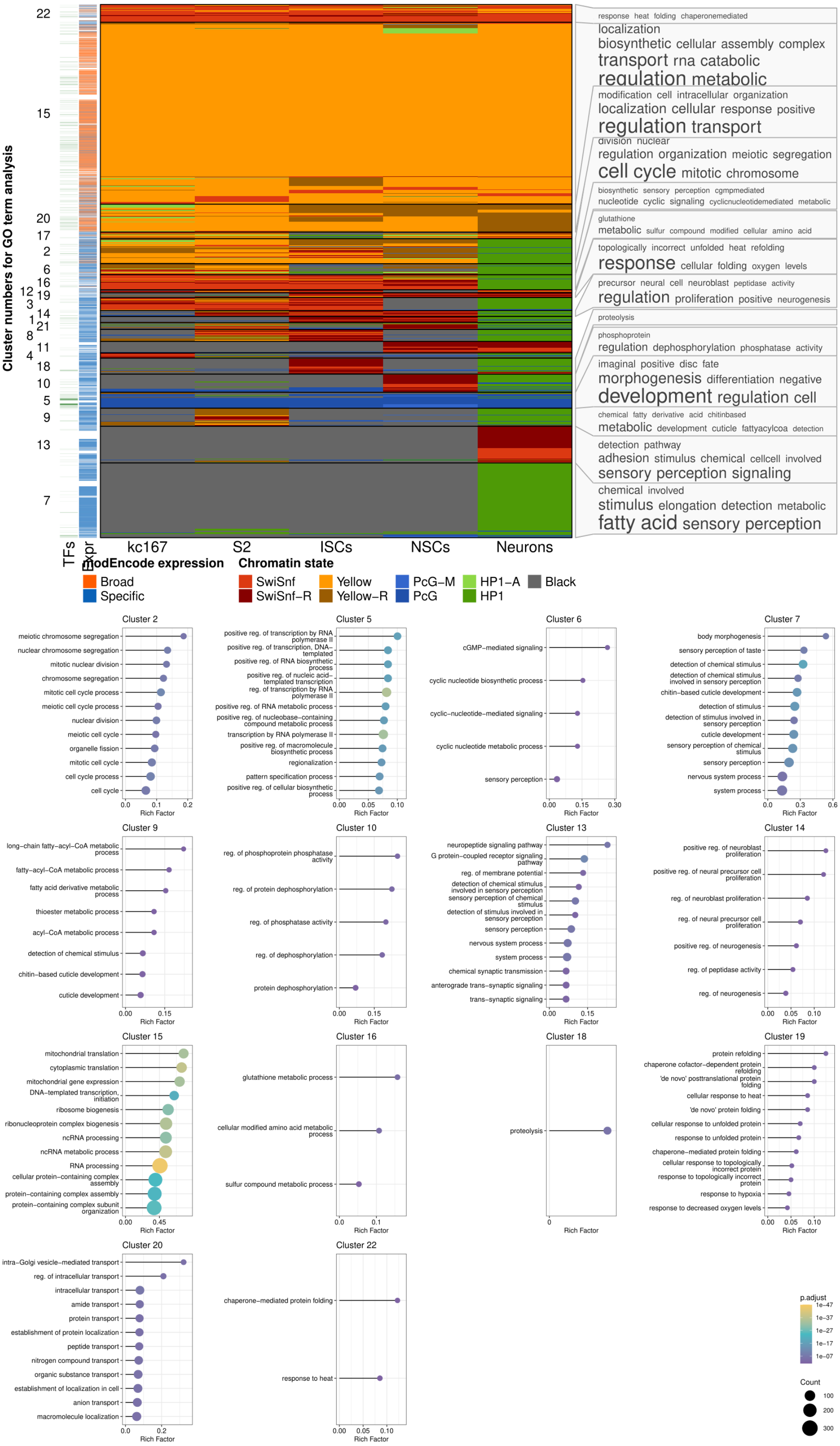

**Figure S14:** Details of GO term enrichment analysis by cluster from Fig. 4.

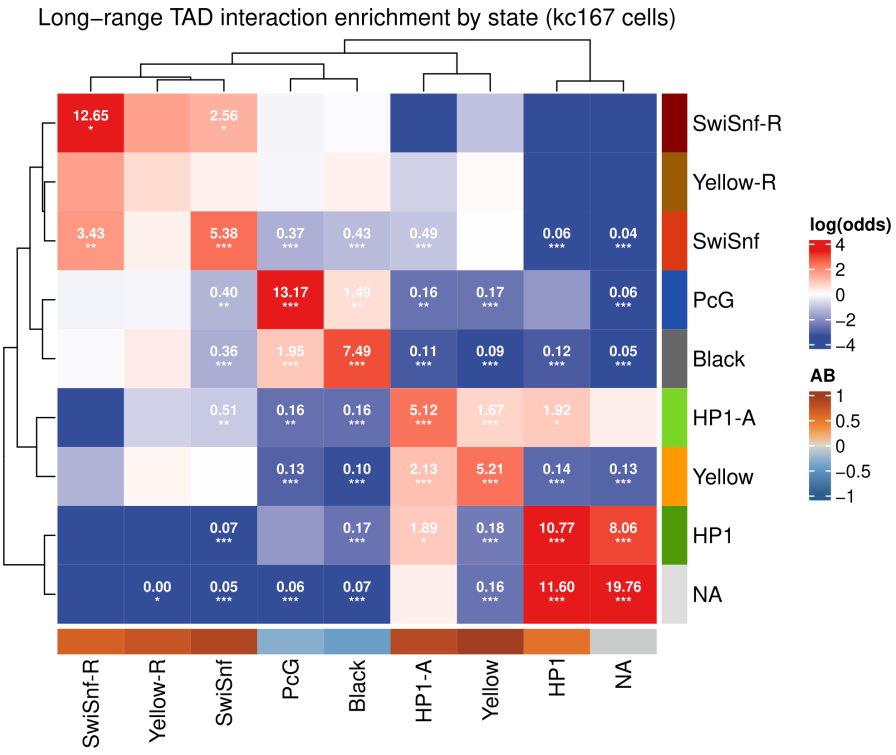

**Figure S15:** Long-range TAD interactions, with heterochromatic TADs separated. The kc167 dataset, including protein binding profiles derived from euchromatic microarray data, is depleted for heterochromatic pericentric regions. As a result, pericentric TADs cannot be directly assigned a chromatin state, and are shown as “NA” in this plot. The Fisher exact test enrichment scores show that these regions are indistinguishable from HP1 TADs in euchromatic arms, allowing us to assign these TADs as HP1 in the analysis in Fig. 6. We note, however, the difference in A/B compartment scores between the pericentric and non-pericentric TADS, with pericentric HP1 TADs being equally likely to be found within A or B compartments.
